# A Key Piece of the Puzzle: The central tetramer of the *Saccharomyces cerevisiae* septin protofilament and Its Implications for Self-Assembly

**DOI:** 10.1101/2023.04.22.537027

**Authors:** Rafael Marques da Silva, Giovanna Christe dos Reis Saladino, Diego Antonio Leonardo, Humberto D’Muniz Pereira, Susana Andréa Sculaccio, Ana Paula Ulian Araujo, Richard Charles Garratt

## Abstract

Septins, often described as the fourth component of the cytoskeleton, are structural proteins found in a vast variety of living beings. They are related to small GTPases and thus, generally, present GTPase activity which may play an important (although incompletely understood) role in their organization and function. Septins polymerase into long non-polar filaments, in which each subunit interacts with two others by alternating interfaces, NC and G. In *Saccharomyces cerevisiae* four septins are organized in the following manner, [Cdc11-Cdc12-Cdc3-Cdc10- Cdc10-Cdc3-Cdc12-Cdc11]_n_ in order to form filaments. Although septins were originally discovered in yeast and much is known regarding their biochemistry and function, only limited structural information about them is currently available. Here we present crystal structures of Cdc3/Cdc10 which provide the first view of the physiological interfaces formed by yeast septins. The G-interface has properties which place it in between that formed by SEPT2/SEPT6 and SEPT7/SEPT3 in human filaments. Switch I from Cdc10 contributes significantly to the interface, whereas in Cdc3 it is largely disorded. However, the significant negative charge density of the latter suggests it may have a unique role. At the NC-interface, we describe an elegant means by which the sidechain of a glutamine from helix α_0_ imitates a peptide group in order to retain hydrogen-bond continuity at the kink between helices α_5_ and α_6_ in the neighbouring subunit, thereby justifying the conservation of the helical distortion. Its absence from Cdc11, along with this structure’s other unusual features are critically discussed by comparison with Cdc3 and Cdc10.

**GRAPHICAL ABSTRACT:** 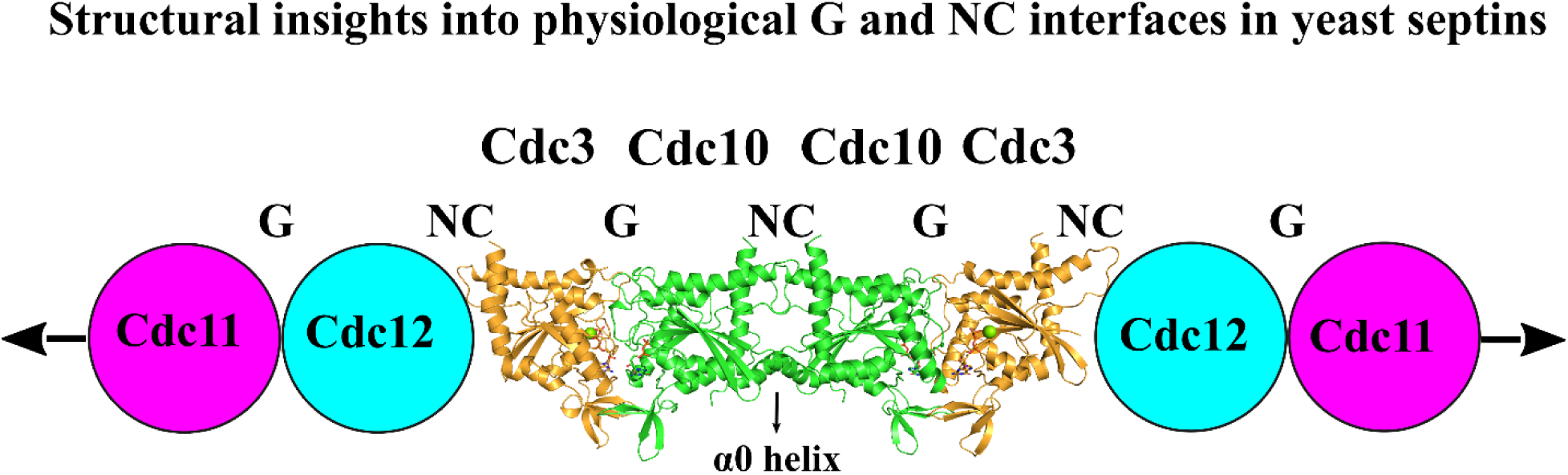

**HIGHLIGHTS:**

- The first crystal structure of a yeast septin heterodimer (Cdc3-Cdc10) provides important insights into their structural biology.
- Identification of common features and differences between yeast and human septins, sheds light on the unique characteristics of yeast septin filaments.
- The Cdc3G-Cdc10_Δ1-10_ crystal structure could be a crucial piece of the puzzle towards obtaining a high-resolution cryo-EM structure of the yeast septin octamer.

## INTRODUCTION

Septins are structural proteins considered to be part of the cytoskeleton and found in a myriad of species, from protozoa to metazoa, but notably absent from plants (Pan et al., 2007; Ryuichi Nishihama, Masayuki Onishi, 2011; Shuman and Momany, 2022). Inside the cell, they polymerize into non-polar filaments in order to accomplish their functions, such as in cell division, creation of diffusion barriers between different intracellular compartments, support for other elements of the cytoskeleton and in anchoring protein binding partners, amongst others (Bridges et al., 2016; Mostowy and Cossart, 2012; Oh and Bi, 2011; Spiliotis and McMurray, 2020).

Depending on the species, the filament can show distinct arrangements, since the number of different septins involved can vary. In mammals, there are 13 different septins clustered into 4 subgroups. A representative from each subgroup (or alternatively just three of them) are combined to form a linear octameric (or hexameric) particle (also called a protofilament). This is the basic unit of end-to-end polymerization (for review see Cavini et al., 2021). In the model organism *Saccharomyces cerevisiae,* there are 7 septins that are related to cell division and sporulation (Versele and Thorner, 2005; Weirich et al., 2008). However, most studies to date have focused on the septins Cdc3, Cdc10, Cdc11 and Cdc12, which form what is known as the canonical protofilament in yeast. These proteins are arranged in the following order in the physiological octamer: Cdc11-Cdc12-Cdc3-Cdc10-Cdc10-Cdc3-Cdc12-Cdc11 (Bertin et al., 2008), where a terminal copy of Cdc11 can interact with another from the next octamer, resulting in the extension of the septin filament. Alternatively, there is the possibility that Cdc11 is replaced by its paralog, Shs1, at the terminal position (Bertin et al., 2008; McMurray et al., 2011), which allows for lateral interaction of the complexes forming arcs, spirals, and rings (Garcia III et al., 2011). On the other hand, Shs1-capped hetero-octamers are not capable of end-to-end association (Booth et al., 2015).

The basic unit of a septin protofilament is the GTP-binding domain (G-domain) of the individual septin monomers. This is dominated by a three-layered αβα sandwich fold, as seen in small GTPases such as Ras (Cavini et al., 2021). Added to this, there is a ∼50 residues C-terminal extension found only in septins (the septin unique element, SUE) (Versele et al., 2004). Besides the G-domain, septins typically contain two further structural domains, known as the variable N-terminal domain (or N-terminal extension, NTE) and the C-terminal domain (or C-terminal extension, CTE). The former is united to the G-domain by a polybasic motif (PB1) believed to be important for membrane association (Bertin et al., 2010; Zhang et al., 1999) and the latter contains pseudo-repetitive sequences characteristic of coiled-coil forming regions (Versele et al., 2004). The G-domain presents GTP-binding and hydrolysis motifs, known as the P-loop (G1), switch I and switch II (G3) regions, G4 (the AKAD motif), and a conserved glycine, a remnant of the G5 motif. Polymerization of the monomers requires that each form two different types of interface with its immediate neighbors on either side. These are called NC and G and alternate along the filament. The SUE, at the end of the G-domain, forms part of both interfaces which explains why septins are able to polymerize while most other small GTPases are not (Cavini et al., 2021).

The G-interface involves the binding pocket where GDP or GTP is buried between the two subunits. On the other hand, the NC-interface is characterized by interactions which involve the N and C termini of each subunit including the PB1 region (between the N- and G-domains) and the coiled coils formed by the C-terminal extensions (Cavini et al., 2021; Mendonça et al., 2021; Sirajuddin et al., 2007). Recently, several papers have described in detail how specific amino acids in human and fly septins determine the necessary specificity to guarantee the interaction of each subunit with its correct partner at each of these two interfaces (de Freitas Fernandes et al., 2022; Mendonça et al., 2021; Rosa et al., 2020; Sala et al., 2016). However, similar information concerning the specific interactions which govern the correct assembly of the yeast octamer is currently lacking even though the order in which the four canonical septins (Cdc3, Cdc10, Cdc11 and Cdc12) associate with one another to generate the full particle appears to have been established (Weems and McMurray, 2017).

Interestingly, the predominant behavior of septins in cells does not seem to involve monomers, but almost certainly these rod-shaped hetero-oligomeric complexes (octamers in the case of yeast). It has already been shown that these rods assemble into short filaments on the plasma membrane, which serves as a platform to generate sufficient local concentration and to reduce the search dimensionality leading to long filament formation as well as to more complex architectures (Bridges et al., 2014; Taveneau et al., 2020). These results show a critical role played by the membrane in the assembly process of septin filaments, but also revive the question of how the septin-membrane interaction actually occurs.

Thus far, yeast and human septins have been the most studied, either due to their participation in basal cellular processes (serving as a model for cell division in Eukarya for example) or because they are related to various human diseases (Fung et al., 2014; Hall and Russell, 2012; Henzi et al., 2021; Van Ngo and Mostowy, 2019). However, since Hartwell’s seminal and inspiring work identifying yeast septins for the first time (Hartwell, 1971), very little progress has been made when it comes to their molecular structure. To date, there is only one yeast septin structure deposited in the Protein Data Bank, Cdc11, which differs from other septin structures regarding certain important aspects of their secondary structure (Brausemann et al., 2016). As a result, almost the entirety of our current knowledge of septin structure comes from humans. In order to fill this gap, we have embarked upon a systematic investigation of the three-dimensional structures of yeast septins, focusing initially on the Cdc3-Cdc10 heterodimer. The structure provides molecular details for two of the interfaces present within the octamer: the G interface between Cdc3 and Cdc10, and the central NC interface formed between two copies of Cdc10, thereby yielding a high-resolution image of the central half of the octameric particle. The structures of both Cdc3 and Cdc10 resemble those found in human septins and differ somewhat from that described for the yeast septin, Cdc11. The new information provided here may be of particular use in understanding septins in general since drawing inferences about other species based on human structures alone can be dangerous, especially given the complexity of septin evolution and the difficulty in correctly assigning orthologs.

## MATERIALS AND METHODS

### Cloning, expression and purification

The DNA coding sequences (CDS) for Cdc3G and Cdc10G GTPase domains, and also for Cdc10 construct, which lacked part of the N-terminal domain but included helix α_0_ and its polybasic region (PB1) (hereafter called Cdc10_Δ1-10_), were codon optimized for *Escherichia coli* expression. DNA sequences were purchased from GenScript (Cdc3G, Cdc10G) and Synbio Technologies™ (Cdc10_Δ1-10_). Additional information about the constructs is detailed in Table S1. The vector pET-15b(+) was used when the G-domains were expressed in isolation, without their septin partner. The septin complexes were co-expressed using the pET-Duet-1 vector (Merck).

*E. coli* BL21 Rosetta™(DE3) cells were used as host for expression. Cells were grown in flasks containing Lysogenic Broth (LB) medium supplemented with chloramphenicol (34 μg·ml-1) and ampicillin (50 µg·ml-1). Cells were grown under constant shaking (150 rpm) at 37 °C. When the culture OD_600nm_ reached around 0.5 – 0.6, protein expression was induced by 0.3 mM (for individual septins) or 0.6 mM (for co-expression) isopropyl 1-thio-β-D galactopyranoside (IPTG) for 16 h, while shaking at 18 °C. Subsequently, the cells were collected by centrifugation (10,000 g for 40 min, at 4°C), and suspended in ice-cold lysis buffer (150 mM NaCl, 50 mM L-Arginine, 50 mM L-Glutamic acid, 5 mM MgCl2, 5 mM β-mercaptoethanol, 10% (v/v) glycerol and 25 mM HEPES pH 8.0). Cell lysis was performed by sonication and the supernatant was collected by centrifugation at 16,000 g for 45 min at 4 °C. The resulting supernatant was then loaded onto a previously equilibrated column containing 2 ml Ni-NTA Agarose (Quiagen^TM^). The column was washed with 10 vol. of lysis buffer and the target proteins were eluted with 2.5 vol. of lysis buffer containing 200 mM imidazole. Size Exclusion Chromatography (SEC) was performed using a Superdex 200 16/70 or Superdex 200 10/300 GL column (GE Healthcare) pre-equilibrated with lysis buffer. Proteins were concentrated using an Amicon^Tm^ Ultra (molecular weight cut-off 30 kDa and 50 kDa) centrifugal filter device (Merck Millipore, Darmstadt, Germany). Fractions from all purification steps were collected and analyzed by SDS-PAGE.

### Size exclusion chromatography coupled with multi-angle light scattering

Size exclusion chromatography coupled with multi-angle light scattering (SEC-MALS) was used to assess the oligomeric state of individual septins and their heterocomplexes (Wyatt Technology, Santa Barbara, CA, USA). Firstly, for the separation of different oligomeric states, 50 μL of each sample (at a concentration of 2 mg·ml−1) were loaded onto a previously equilibrated Superdex 200 10/300 GL column (20 mM Tris-HCl and 150 mM NaCl buffer (pH 7.8) coupled to an HPLC (Waters, Milford, MA, USA). Secondly, light scattering data from the samples were recorded by a three-angle light scattering detector; mini DAWN® TREOS® (Wyatt Technology, Santa Barbara, CA, USA). Finally, for data collection and analysis the software Wyatt ASTRA 7 (Wyatt Technology Corporation, Santa Barbara, CA, USA) was used.

### Nucleotide identification

The presence (or absence) of endogenous GTP or GDP in the Cdc3G and Cdc3G-Cdc10G samples was verified as described by (Seckler et al., 1990), with minor modifications. 500 μL of each sample at a concentration of 25 µM were incubated with 200 μL of cold HClO_4_ (4 °C) for 10 mins and then centrifuged at 16,000 g for 10 min, also at 4 °C, for protein precipitation. Subsequently, 600 μL of the supernatant was transferred to a new microtube and then neutralized with cooled solutions of KOH 3 M (100 μL), K_2_HPO_4_ 1 M (100 μL) and 5 M acetic acid (80 μL). After incubation for 18 hours at -20 °C, samples were centrifuged once more at 16,000 g for 10 min, at 4 °C. An ion exchange column (Waters^TM^ Protein Pack DEAE-5 PW 7.5 mm x 7.5 cm) coupled to a HPLC Alliance 2695 chromatography system, previously equilibrated in 25 mM Tris-HCl pH 8.0, was used for nucleotide detection. 200 μL of each sample were loaded onto the column and the nucleotides were eluted by a linear NaCl gradient (0.1 – 0.45 M in 10 min.). 200 µL samples of buffer containing only GDP or GTP at 50 µM were run for comparison purposes. Absorbance was monitored at 253 nm.

### Crystallization, data collection and structure determination

Septin complexes of Cdc3G-Cdc10G and Cdc3G-Cdc10_Δ1-10_ were crystallized by the sitting drop vapour diffusion method using the PACT++ HTS screening kit (Jena Biosciece). 0.6 μl of the samples at a concentration of 13 mg.mL^-1^ was mixed with 0.3 μl of 20 % w/v polyethylene glycol 3,350, 200 mM sodium formate and 100 mM BIS-TRIS propane pH 6.5 (Cdc3G-Cdc10G) or 27% polyethylene glycol 3,350, 100 mM BIS-TRIS propane pH 6,5 and 270 mM sodium iodide (Cdc3G-Cdc10_Δ1-10_). Crystals appeared after 24 hours of incubation at 20°C and were harvested after five days. Polyethylene glycol 200 was used as a cryoprotectant and crystals were kept in liquid nitrogen until data collection. X-ray diffraction data was collected at 100 K on the Manacá beamline of the Sirius Synchrotron (Campinas, Brazil). Indexation, integration and data reduction/scaling was done by *autoPROC* (Vonrhein et al., 2018). Molecular replacement was carried out by Phaser (McCoy et al., 2007), using a Cdc3-Cdc10 *AlphaFold2* model as a search model (Jumper et al., 2021; Mirdita et al., 2022). Alternate rounds of refinement and model building were conducted using *phenix.refine* (Adams et al., 2010) and Coot (Emsley and Cowtan, 2004). DIMPLOT (Laskowski and Swindells, 2011) was used to identify the interacting residues across the G and NC-interfaces and the APBS-PDB2PQR software suite was used to calculate the electrostatic potential over the dimer surfaces employing default parameters (Jurrus et al., 2018). Pymol v2.05 was used for image generation. Data collection, refinement statistics and PDB code are summarized in Table S2.

## RESULTS and DISCUSSION

### Protein purification and oligomeric state characterization

Recombinant expression assays of the full-length versions of Cdc3 and Cdc10 have shown high instability and insolubility (Baur et al., 2019; Farkasovsky et al., 2005). Only a truncated version of Cdc3, including only the G-domain has been previously purified in a soluble state (Baur et al., 2019). For the present study we initially attempted to recombinantly express Cdc3G and Cdc10G separately. Soluble Cdc3G was purified by affinity (IMAC) and size exclusion (SEC) chromatography (Fig. S1A), showing it to be monomeric in solution as revealed by SEC coupled with multi-angle light scattering (SEC-MALS) (Fig. 1). However, we were unable to purify Cdc10G due to its recalcitrant insolubility, as can be seen on SDS-PAGE where it appears exclusively in the pellet fraction (P) (Fig. S2A). The identity of both proteins was confirmed by mass spectrometry analysis (Fig. S2B, C).

**Figure 1.**
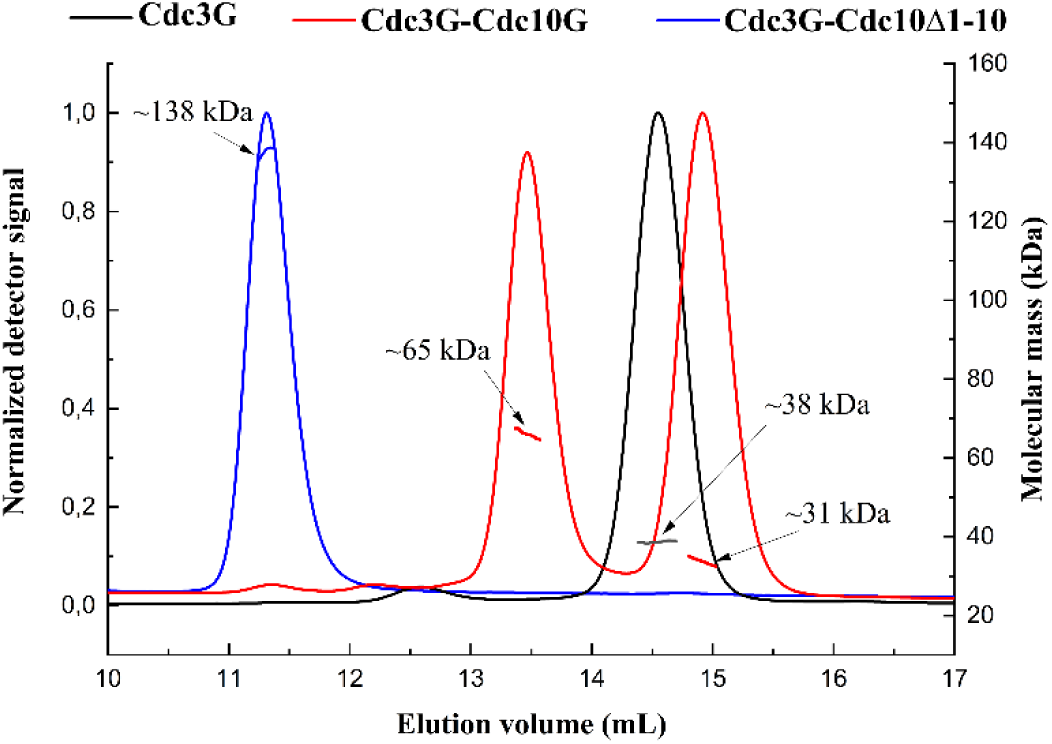
SEC-MALS profiles show the oligomeric state of Cdc3G (∼38 kDa corresponding to a monomer), Cdc3G-Cdc10G (∼31 and ∼65 kDa corresponding to a mixture of monomers and dimers, respectively) and Cdc3G-Cdc10_Δ1-10_ (tetramers with a mass of ∼138 kDa).

According to the literature, the presence of the correct partner can make a significant difference to the solubility and stability of individual septins (Baur et al., 2019; Kumagai et al., 2019). Following the yeast septin model of the octameric protofilament (Bertin et al., 2008), we would expect a complex formed between Cdc3G and Cdc10G employing a G-interface. Thus, as an attempt to purify Cdc10G, the coding sequence of both Cdc3G and Cdc10G were inserted into the pET-Duet vector for co-expression. The presence of two bands on SDS-PAGE (Fig. S2A) confirms the co-purification of Cdc3G-Cdc10G, demonstrating the positive influence of the correct G-interface partner (Cdc3) on the stability and solubility of Cdc10 (as has also been observed for their full-length versions (Baur et al., 2019)). In addition, the absence of the N- and C-terminal domains suggests that complex formation via the G-interface is viable as reported for septins from other organisms (de Freitas Fernandes et al., 2022; Rosa et al., 2020). The SEC profile shows two oligomeric states in solution (Fig. S1B), which were confirmed to be dimers and monomers by SEC-MALS (Fig. 1).

Like the G-interface, the NC-interface is also essential for the formation of septin filaments (Bertin et al., 2010; Sirajuddin et al., 2007; Weems and McMurray, 2017). In *S. cerevisiae*, the NC-interface between two Cdc10 subunits at its center is integral to the formation of canonical octamers (Cdc11-Cdc12-Cdc3-Cdc10-Cdc10-Cdc3-Cdc12-Cdc11) and previous studies have emphasized that a structural element at the N-terminus of the G-domain (helix α_0_) is critical for stabilizing this interaction (Bertin et al., 2010). Furthermore, it is α_0_ that harbours a polybasic region (PB1) important for membrane association, a process in which Cdc10 appears to be intimately involved (Bertin et al., 2010). For several reasons, therefore, the α_0_-helix from Cdc10 is of particular interest and so a new construct including it was elaborated (named Cdc10_Δ1-10_ as it lacked only the first ten residues of the full protein) and this was co-expressed and co-purified with Cdc3G to assess if this affected the oligomeric state of the complex. Analysis of the SEC-MALS profile shows that Cdc3G-Cdc10_Δ1-10_ forms stable tetramers in solution (Fig. 1) as compared with only dimers at most for Cdc3G-Cdc10G, suggesting the formation of a physiological Cdc10-Cdc10 NC-interface facilitated by the presence of the α_0_-helix. Bertin et al., 2010 demonstrated by negative stain TEM that by removing the α_0_-helix from Cdc10, the formation of octamers and filaments and their association with membranes is compromised (observing instead mostly tetramers). This emphasizes the role of the Cdc10 α_0_-helix in maintaining the integrity of octamers (by stabilizing the central Cdc10-Cdc10 NC interface) and thereby the formation of heterofilaments and their association with biological membranes. Indeed, α_0_ from Cdc10 appears to play a dominant role over other septins within the complex (Bertin et al., 2010). For this reason, we decided to determine the crystal structures of both the Cdc3G-Cdc10G and Cdc3G-Cdc10_Δ1-10_ complexes.

Nucleotide binding and hydrolysis has been shown to be a pivotal event in complex and filament formation of yeast septins (Weems and McMurray, 2017). By detecting the presence and nature of the nucleotide released after protein denaturation, we observed that Cdc3G when expressed in isolation is purified free of nucleotide (Fig. S1C). The apo form of Cdc3 has been previously reported, where it was also shown it to be incapable of binding and hydrolyzing exogenous nucleotides (Baur et al., 2019). The authors suggest that the substitution of the catalytic threonine of switch I (ThrSw1 nomenclature due to Cavini et al, 2021) by lysine, is responsible for the absence of the “loaded spring” mechanism for hydrolysis. This behavior is observed in human septins of the SEPT6 subgroup, which also lack ThrSw1 and are purified as nucleotide-free monomers which are catalytically inactive (Macedo et al., 2013; Zent and Wittinghofer, 2014). GDP/GTP uptake assays of full-length Cdc3-Cdc10 showed that the complex takes up GDP but not GTP, and there was no prior detection of the presence of bound nucleotides (Baur et al., 2019). In contrast, we identified both GTP and GDP bound to the G-domain dimer, Cdc3G-Cdc10G, a result not unexpected due to the presence of a catalytically inactive (Cdc3) and a catalytically active (Cdc10) septin (Fig. S1C). The bound nucleotides were subsequently confirmed in the crystal structures described below.

### The crystal structures of Cdc3G-Cdc10G and Cdc3G-Cdc10***_Δ_***_1-10_

Although yeast septins have been widely studied in terms of their biochemistry, cell-biology and physiology of filament formation, structural studies reported to date have been limited to only the structure of Cdc11G at medium resolution (Brausemann et al., 2016). Using a “divide and conquer” strategy, which is based on studying each of the interfaces of the octamer in isolation, we solved the crystal structures of the heterodimeric complexes formed by Cdc3G-Cdc10G and Cdc3G-Cdc10_Δ1-10_, the first from yeast to be described.

#### Overall description and interface arrangements within the crystal

The structure of Cdc3G-Cdc10G and Cdc3G-Cdc10_Δ1-10_, at 2.22 Å and 2.66 Å resolution respectively, were solved by molecular replacement using as a search model the Cdc3-Cdc10 dimer generated by AlphaFold2 (Jumper et al., 2021; Mirdita et al., 2022). All data collection and quality control parameters are within expected values (Table S2). With a classic septin fold based on a three layered αβα sandwich (Fig. 2) (Cavini et al., 2021), we observed that Cdc3G, Cdc10G and Cdc10_Δ1-10_ are all very similar to other septin structures already described in other species (de Freitas Fernandes et al., 2022; Mendonça et al., 2021; Rosa et al., 2020), reinforcing that many of the structural characteristics are highly conserved.

**Figure 2.**
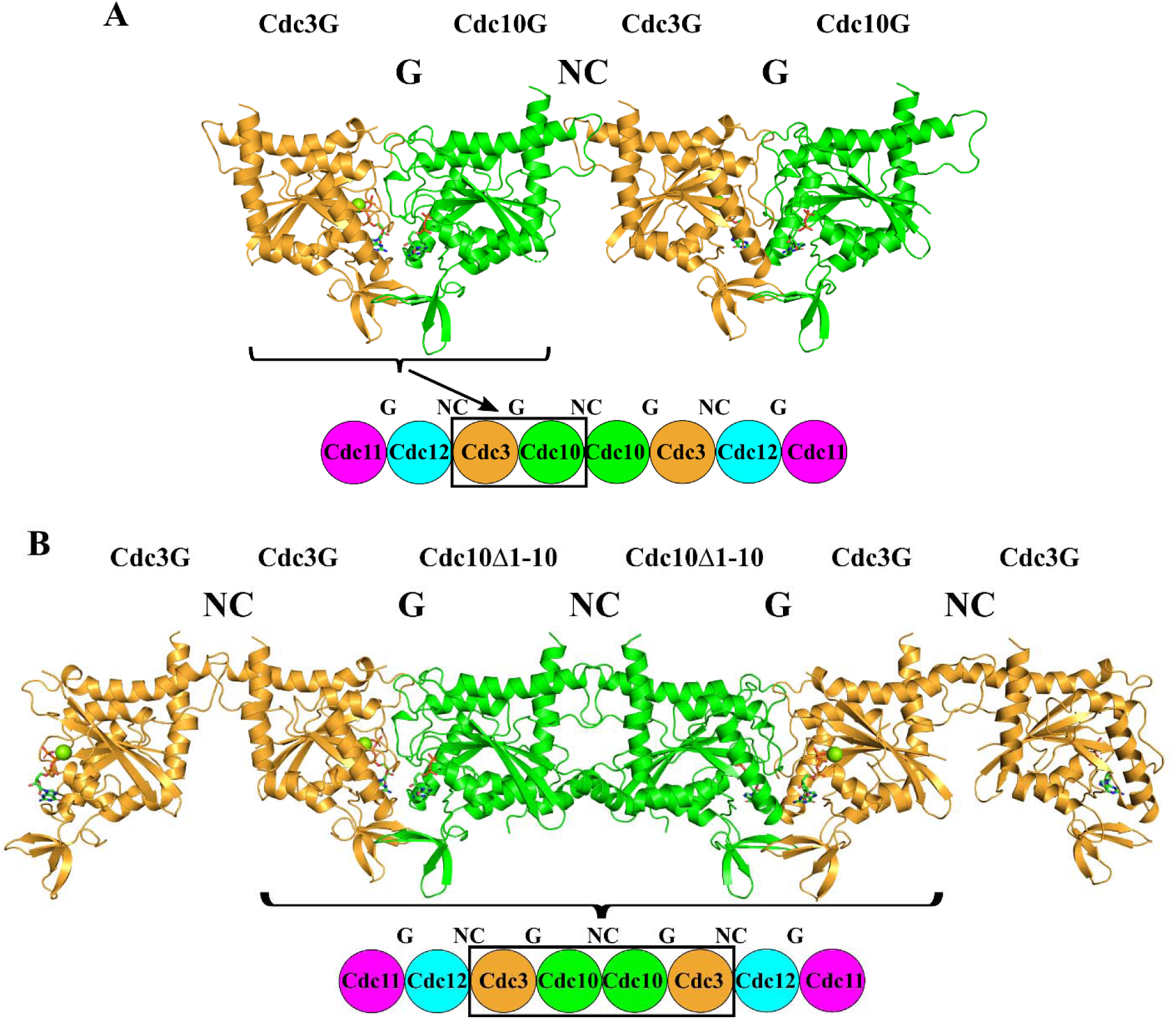
Crystal structures of Cdc3G-Cdc10G and Cdc3G-Cdc10_Δ1-10_ reveal the classic αβα sandwich fold of septins, with GTP+Mg^2+^ and GDP bound to Cdc3 and Cdc10, respectively. A) The Cdc3G-Cdc10G G-dimer forms filaments in the crystal using the non-physiological NC-interface between Cdc10G and Cdc3G. B) The inclusion of the α_0_-helix in Cdc10_Δ1-10_ allows for the formation of filaments in the crystal with physiological G-interfaces (Cdc3G-Cdc10_Δ1-10_) and two types of NC-interfaces, one physiological (Cdc10_Δ1-10_-Cdc10_Δ1-10_) and one promiscuous (Cdc3G-Cdc3G).

The Cdc3G-Cdc10G structure has a G-interface dimer in the asymmetric unit, with each subunit bound to a nucleotide: Cdc3 bound to Mg^2+^/GTP (as expected for a catalytically inactive septin) and Cdc10 to GDP (a catalytically active septin) (Fig. S3). This is consistent with the nucleotide content described above. Besides the physiological G-interface, in the crystal filaments are formed using a non-physiological (promiscuous) NC-interface between Cdc3 and Cdc10 rather than a physiological interface using two copies of Cdc10 (as shown in the schematic model in Fig. 2A). Non-physiological NC-interfaces have been observed in crystal structures of heterodimers from other species, demonstrating that the G-domain alone is insufficient to guarantee the formation of the correct NC pairs (de Freitas Fernandes et al., 2022; Rosa et al., 2020). On the other hand, the Cdc3G-Cdc10_Δ1-10_ structure has two dimers in the asymmetric unit (both forming G-interfaces). As in the previous case, Cdc3G and Cdc10_Δ1-10_ are bound to GTP and GDP respectively. However, on this occasion the filaments formed in the crystal are generated by two different types of NC-interface. One is promiscuous (Cdc3-Cdc3) and the other physiological (Cdc10-Cdc10) (Fig. 2B). Although the influence of different crystallization conditions cannot be excluded, these results appear to demonstrate that it is the presence of the α_0_-helix which is critical for the formation of the correct NC interface between the two copies of Cdc10 (described in detail below). Furthermore, it strongly suggests that the tetramers observed in solution (Fig. 1) are formed between two Cdc3-Cdc10 dimers through an NC-interface (Cdc3-Cdc10-Cdc10-Cdc3). This represents half of the octameric particle and resides at its center.

### The ***α***_0_-helix is critical for the physiological NC-interface of Cdc10

The NC-interface observed between two copies of Cdc10_Δ1-10_ is essentially complete. The construct lacks only ten residues from the full protein at its N-terminus and the first residue of the expressed protein (Ser11) is disordered in the crystal structure. Although it was anticipated that the short C-terminal domain (303-322) might make a significant contribution to the stability of the interface, in fact it is completely disordered beyond Ile301, at the end of α_6_. The interface is observed to be in the “open” conformation (Castro et al., 2020), displays a large internal cavity as described previously (Mendonça et al., 2021) and shows the classical network of salt bridges. These have been reported to form the upper part of the interface and have been extensively described previously (Castro et al., 2020; Mendonça et al., 2021) and will not be discussed further here. Similarly, we will not go into detail about the homodimeric NC-interface involving Cdc3, which appears in the crystal structure of Cdc3G-Cdc10_Δ1-10_, but is considered to be promiscuous, i.e. not expected to exist in physiological septin filaments. The most interesting feature of the Cdc10 interface is the presence of the two α_0_ helices which line the floor of the NC-cavity.

The two α_0_ helices lie tucked within the NC-interface where they lie antiparallel to one another in a similar fashion to that seen in the small number of other septin structures where they are also present (Fig. 3A) (Mendonça et al., 2021; Sirajuddin et al., 2007). The helices make symmetrical VDW contacts via the sidechains of Ile22 and Leu26. These would be lost if Cdc3 were to pair with Cdc10 across the NC-interface as observed in the Cdc3G-Cdc10G crystal structure. This is due to substitution of Leu26 by Ser113 in Cdc3. As mentioned above, it therefore appears that it is the presence of the α_0_ helix and the additional contacts that it provides which favor the physiological interface observed in the crystal structure of Cdc3G-Cdc10_Δ1-10_. This is entirely consistent with the observations made by Bertin et al., (2010) using negative stain TEM in which the use of a construct lacking the greater part of α_0_ resulted in destabilization of the NC interface to the point of producing only tetramers at high salt concentration (presumably Cdc11-Cdc12- Cdc3-Cdc10) or inverted octamers at low salt concentration (Cdc10-Cdc3-Cdc12-Cdc11-Cdc11- Cdc12-Cdc3-Cdc10). Furthermore, the authors were able to show that octamers containing a Ile22Glu mutant had a significantly less stable central interface and anomalous polymerization behaviour. This is presumably due to the disruption of the hydrophobic contacts made between the two α_0_ helices.

**Figure 3.**
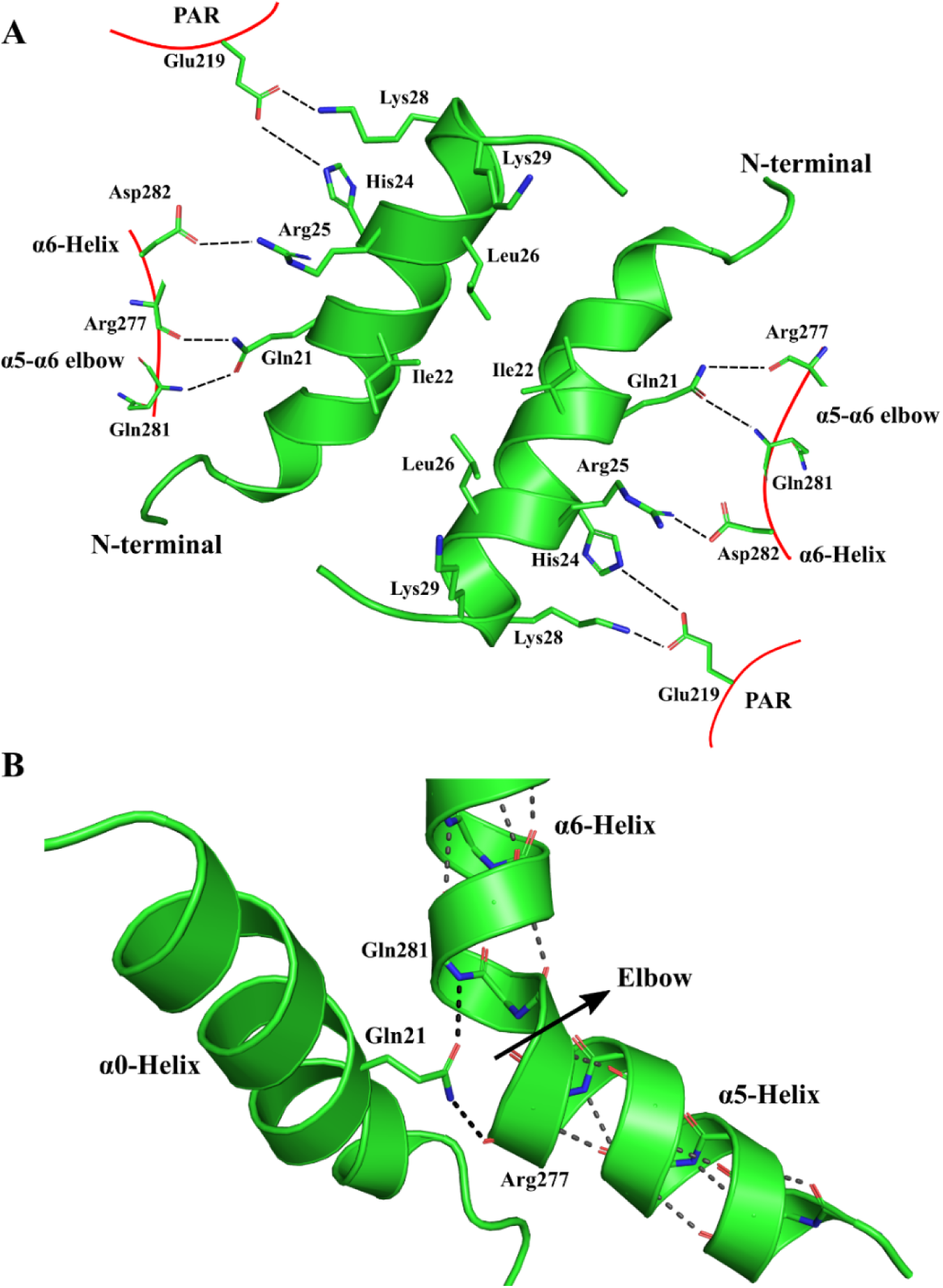
Key aspects of the physiological Cdc10_Δ1-10_-Cdc10_Δ1-10_ NC-interface. A) Interaction of the α_0_- helix with the polyacidic region (PAR) and helix α_6_. At its C-terminus, α_0_has the polybasic region 1 (PB1) of which His24 and Lys28 interact with Glu29 (from the PAR) and Arg25 with Asp282 of α_6_. Gln21, a largely conserved amino acid in septins, forms hydrogen bonds with the mainchain of Arg277 and Gln281 from the α_5_-α_6_ elbow. The α_0_-helices of both subunits in the NC-interface lie antiparallel, allowing for the formation of hydrophobic contacts between Ile22 and Leu26. B) Interaction between Gln21 from α_0_of one subunit and the α_5_-α_6_ elbow (Arg277 and Gln281) from the other. The hydrogen bond network along the α_5_-α_6_ helices, which is interrupted at the kink (elbow), is re-established by the sidechain of Gln21, which appears to be an integral part of the NC-interface, contributing to its stability.

The C-terminal region of α_0_ corresponds to the polybasic region known as PB1 (Fig. 3A) (Cavini et al., 2021; Zhang et al., 1999). This has been reported to be important for binding to biological membranes, particularly those containing phosphotidylinositol-4,5-bisphosphate (PIP2) (Bertin et al., 2010). Indeed, the PB1 region of Cdc10 appears to be of particular importance for the association between yeast septin filaments and PIP2 containing artificial membranes since several lines of investigation show it to present a markedly more pronounced effect when compared to Cdc3, Cdc12 or Cdc11. Details of the structure of PB1 have been described previously, based on the heterodimeric interface formed between human SEPT6 and SEPT7 in the hexameric complex composed of SEPT2, SEPT6 and SEPT7 (Mendonça et al., 2021). The most striking observation, which is repeated here for Cdc10, is the fact that the basic residues themselves are occupied in ionic interactions and orientated such that their involvement in membrane interactions appears impossible (Fig. 3A). This led to the conclusion that the interface must suffer a conformational change in order to expose PB1. Such plasticity has been observed at the NC-interface between members of the SEPT3 group of human septins, a dimer of which occupies the central position of human octamers, analogous to Cdc10 in yeast.

Together, the observations from both human and yeast models suggest that the NC-interface which resides at the center of the octamer may present unique properties. Particularly, squeezing the interface may provide a mechanism for exposing PB1 and enhancing membrane interaction. This represents an interesting working hypothesis, but clearly further studies are required on the structural biology of the yeast system before any conclusions can be drawn. Nevertheless, it would be consistent with the special role observed for PB1 in Cdc10 when compared to the remaining components of the octamer.

Typically, four basic residues participate in PB1 distributed in the following manner [++xx++], where + is a basic residue and x is variable. In Cdc10 these correspond to [H_24_RLLKK_29_] (Fig. 3A). The interactions described previously for the human complex hold largely true for Cdc10: Arg25 and Lys28 interact with the neighboring subunit by forming salt bridges with Asp282 of α_6_and the polyacidic region (PAR) prior to α_5’_ respectively, while Lys29 points into the NC-cavity (Fig. 3A). His24 appears to play a different role to its human homologue (Arg42 in SEPT7) and, together with Lys28 interacts with Glu219 of the PAR. Arg42 in SEPT7, on the other hand was described to point towards the C-terminus of α_5_, aligning with its electrical dipole. However, with the higher resolution now available it has become clear that the residue which is actually involved in satisfying the H-bonding potential at the C-terminus of α_5_ is Gln21, as described below. Consistent with the interactions made by the basic residues across the interface, when they are simultaneously substituted by alanine, the central interface of the octamer is once again weakened (Bertin et al., 2010).

Helices α_5_ e α_6_ are part of the NC-interface and are traditionally described as separate elements of secondary structure (Cavini et al., 2021; Sirajuddin et al., 2007). However, this is somewhat arbitrary, and it would be equally valid to describe the entire structure as forming a single entity, namely a helix bearing a ∼50° kink. However, the wide kink angle (Wilman et al, 2014) justifies their original denomination as two different elements of secondary structure (Sirajuddin et al, 2007) and we do not propose altering the current nomenclature. The kink is centered on residue His279 in Cdc10 (Wilman et al., 2014) and is conserved in all known structures of septins except for Cdc11 (Brausemann et al., 2016). This is indicative of a possible functional role for the kink as observed in other protein families including G-protein coupled receptors (Law et al., 2016). We have called this structure the α_5_-α_6_elbow (Mendonça et al., 2021).

Kinks (or elbows) in α-helices disrupt the regular pattern of hydrogen bonding leaving mainchain amino groups and carbonyls unsatisfied. In the case of septins, these face towards the NC- cavity and the free backbone amino group of Gln281 (in Cdc10) and the carbonyl of Arg277 form hydrogen bonds with the side chain amide of Gln21 from α_0_ of the neighbouring subunit (Fig. 3B). The sidechain amide emulates a peptide group thereby partially reconstructing the H-bonding pattern of the two helices (Fig 3B). For the glutamine side chain to fit within the elbow, it is necessary that the kink angle be wide, as we observe. Gln21 is conserved in all human and yeast septins except for Cdc11, indicating that it, together with the α_5_-α_6_ elbow, are integral parts of the NC-interface, aiding in ensuring its stability. The importance of Gln21 in establishing a viable NC interface may, in fact, be more important than the basic residues of PB1, which may be retained in the interface for safe keeping when not interacting with membranes.

### Structural relationships between Cdc10 and the human SEPT3 subgroup

Phylogenetic analysis classifies all septins into five different groups, being Cdc10 the only yeast septin which is grouped together with one of the mammalian subgroups (SEPT3) (Pan et al., 2007; Shuman and Momany, 2022). Likewise, comparing the arrangement of the octamer for yeast (Cdc11-Cdc12-Cdc3-Cdc10-Cdc10-Cdc3-Cdc12-Cdc11) and human septins (SEPT2- SEPT6-SEPT7-SEPT3-SEPT3-SEPT7-SEPT6-SEPT2), shows that Cdc10 occupies a position equivalent to SEPT3 at the center of the particle. The two crystal structures reported here for the different constructs of Cdc10 show that it not only shares a close evolutionary relationship and position within the filament with SEPT3, but also preserves structural features that are unique to the subgroup (Fig. 4).

**Figure 4.**
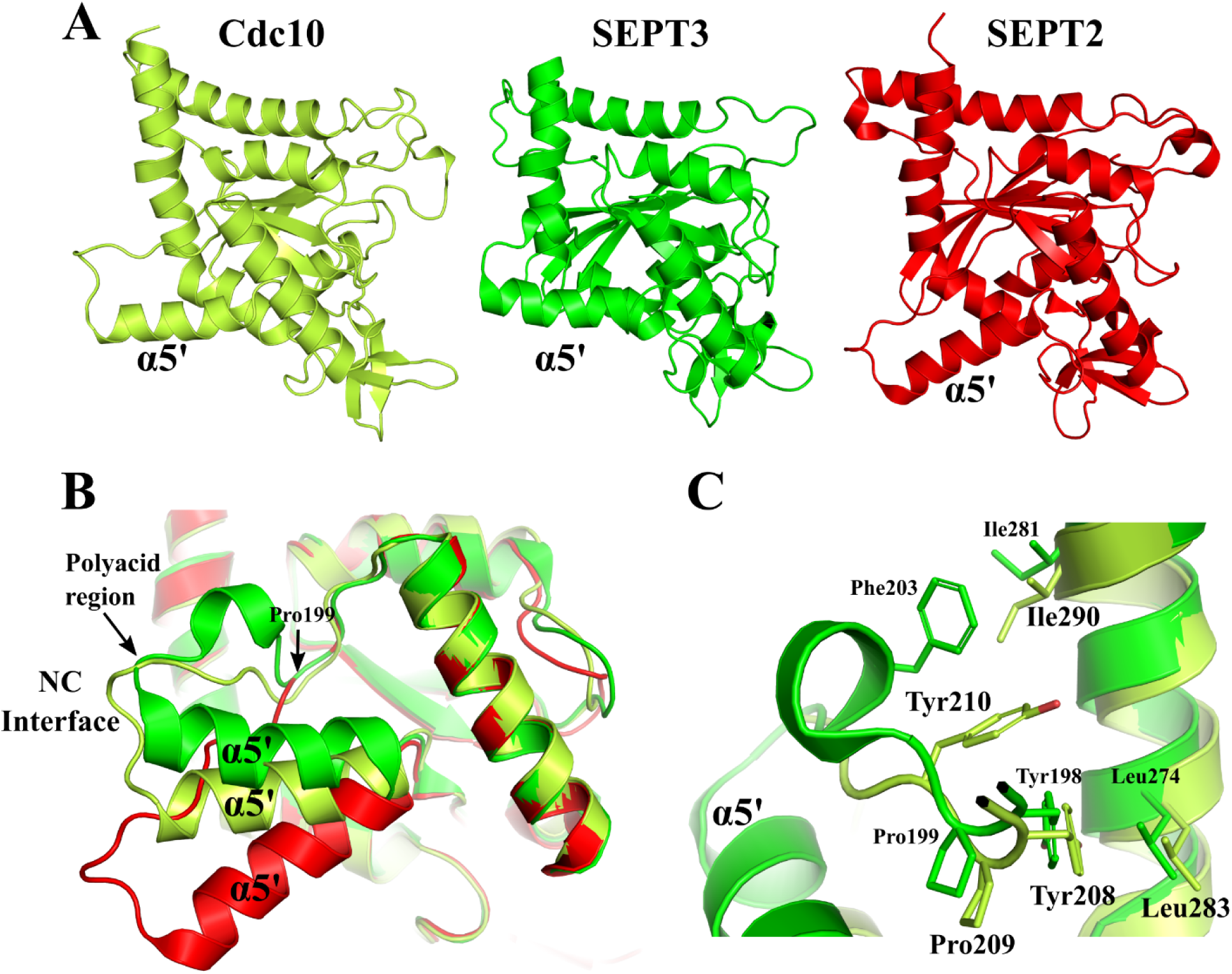
Structural relationships between Cdc10 and human SEPT3. A) Ribbon diagrams for Cdc10, SEPT3 (4Z54) and SEPT2 (6UPQ) show the different orientations for the α_5’_-helix. B) Reorientation of α_5’_ in Cdc10 and SEPT3 in relation to SEPT2, (used as an example to represent the conventional orientation normally observed). Pro199 of SEPT3 is responsible for deviating the course of the polypeptide chain including the loop containing the polyacidic region which becomes raised within the NC-interface. Cdc10 presents Pro209, homologous to Pro199, a characteristic of the SEPT3 subgroup. C) Cdc10 retains the interaction Tyr208(198)→Leu283(274) but the absence of Phe203 (conserved in the SEPT3 subgroup) could be related to a less pronounced dislocation of α_5’_ to that observed in SEPT3.

Human SEPT3 subgroup septins show a dislocation of the α_5’_-helix compared to septins from other subgroups (Castro et al., 2020), a characteristic also observed in Cdc10 (Fig. 4A and 4B). Castro et al., 2020 show the importance of a characteristic proline (Pro199 in Fig. 4B) in altering the course of the main chain of the α_4_-α_5’_loop, which consequently impacts on the orientation of the α_5’_-helix. Cdc10 presents a proline (Pro209) in the same position causing helix dislocation similar to that seen in SEPT3 (Fig. 4B). Other interactions also contribute to this reorientation in SEPT3, including those formed between Tyr198 and Leu274 and a hydrophobic contact between Phe203 (located in the PAR) and Ile281 of the α_6_-helix (Fig. 4C) (Castro et al., 2020). Cdc10 retains a similar Tyr208-Leu280 interaction but the absence of Phe203 is compensated for by Tyr210 (Fig. 4C). This difference could explain why the reorientation of α_5’_ is smaller in Cdc10 with respect to human SEPT2 (a 10Å shift at its N-terminus) as compared with the human SEPT3 subgroup, where the shift can be up to 20Å (Fig. 4B). The consequences of the reorientation of α_5’_ is similar in both cases, namely its N-terminus is lifted within the NC-interface taking the preceding PAR with it.

It has been shown that the polybasic region 2 (PB2), between α_2_ and β_4_, is involved in the opening and closing mechanism of the NC-interface of the SEPT3 subgroup (Castro et al., 2020). PB2 makes significantly more interactions with the PAR of the α_4_-α_5’_ loop of the neighbouring subunit when in the “closed” state compared with the “open” state. Cdc10 preserves in part the PB2 region but with fewer basic amino acids than in the SEPT3 sub-group (2 against 4 consecutive charges found in the latter) (Fig. S4A). Nevertheless, a similar mechanism may be at play. However, we note that the conformation of this region is similar in both Cdc3G-Cdc10G and Cdc3G-Cdc10_Δ1-10_, with the only difference being the orientation of Arg136 (Fig. 4SA) which is able to form a salt bridge between with Glu214 of the PAR even in the “open” conformation seen in the crystal structure. It is, therefore, still far from clear if the open-closed conformational change observed in human septins could be expected to apply to yeast and even less so if it is relevant *in vivo*. Nevertheless, there seems to be considerable evidence suggesting that the central interface of the octamers in both cases may have unique structural properties which merit direct investigation.

### Characteristics of the physiological G-interface

The Cdc3-Cdc10 structures show for the first time a physiological G-interface in yeast septins, allowing comparison of the structural determinants of this interaction with the physiological G-interfaces seen in human septins. Since the Cdc3 construct used here is limited to the G-domain and the heterodimeric interface is intact, our structure is unable to shed light on the details of the predicted interaction in *cis* between the N-terminal domain of Cdc3 and its own G-interface which is believed to control important aspects of oligomer assembly (Weems and McMurray, 2017). For descriptive purposes we will use the structure with the higher resolution (Cdc3G-Cdc10G), and to facilitate the interpretation, we will divide the interface into three sections: upper, middle, and lower regions in accordance with de Freitas Fernandes et al.,2022 (Fig. 5A).

**Figure 5.**
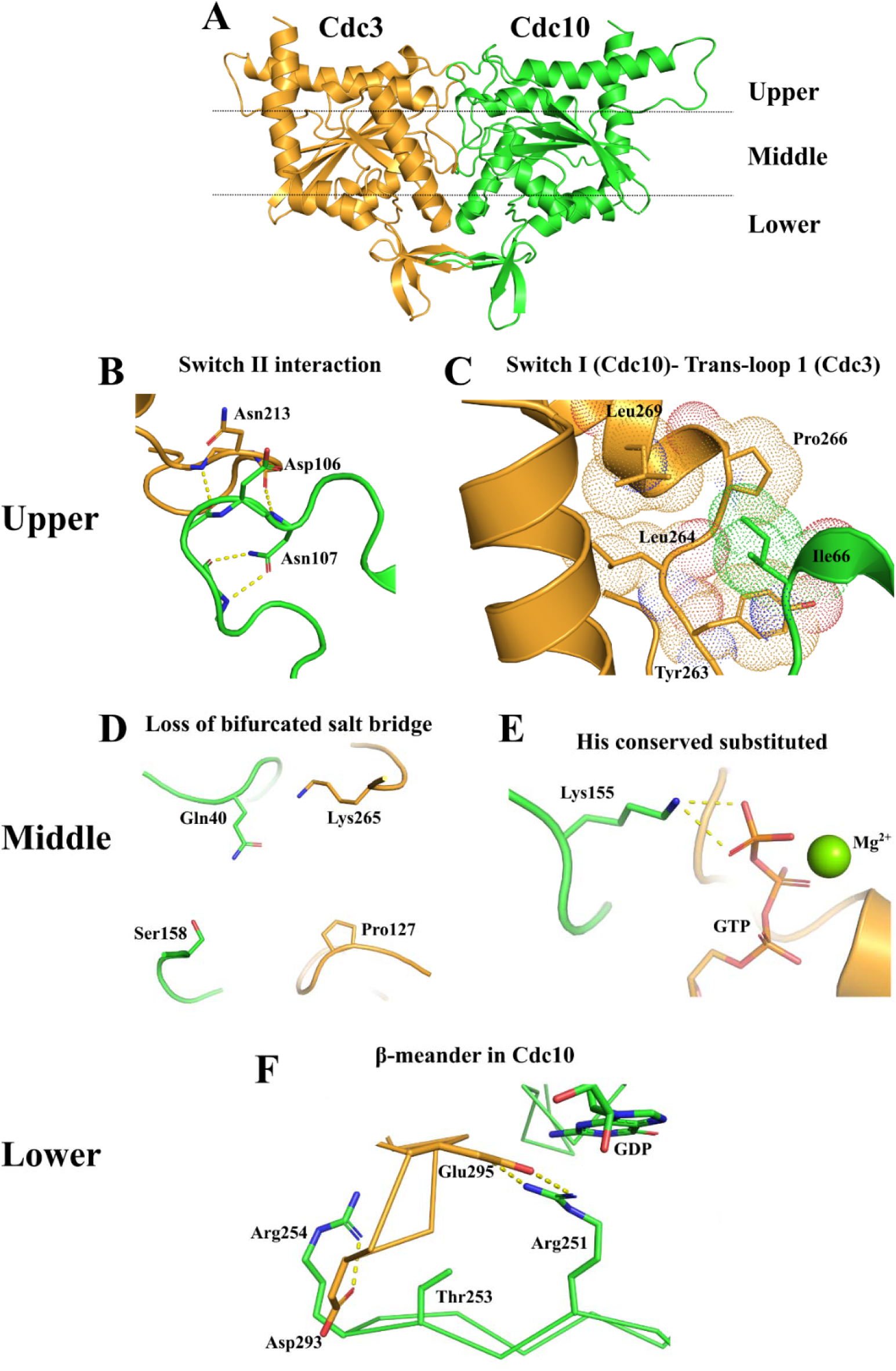
The physiological G-interface in Cdc3G-Cdc10G. A) The dimer was divided into upper, middle, and lower sections for easier description. B) The β-bridge formed by the switch II regions of both subunit is shown. This bridge unites As213 of Cdc3 and Asp106 of Cdc10. C) Ile66 from the switch I of Cdc10 fits into the hydrophobic cavity formed by the trans-loop 1 (Tyr263, Leu264, Pro266, and Leu269) of Cdc3. D) The loss of the bifurcated salt bridge in the G-interface due to substitutions at three of the four residues which define it. E) The substitution of a conserved histidine by Lys155 in Cdc10. Despite the substitution, an interaction GTP is maintained (via its γ phosphate). F) Two salt bridges between Arg251(Cdc10) - Glu295(Cdc3) and Arg254(Cdc10) - Asp293(Cdc3), involving positively charged residues located in the β- meander of Cdc10. Additionally, Thr253 of Cdc10 in indicated. In human septins this threonine is characteristic of the SEPT3 subgroup.

The upper section includes the interaction between the switch II regions of the two subunits. These form the classical β-bridge described for physiological G-interfaces observed in other species (Fig. 5B) (Brognara et al., 2019; de Freitas Fernandes et al.,2022; Rosa et al., 2020) and in the complexes described here is defined by Asn213(Cdc3) and Asp106(Cdc10).

Cdc10 occupies the central position of the octamer in the yeast septin model (Bertin et al., 2008). Therefore, it is expected that the Cdc3-Cdc10 dimer would be analogous to SEPT7- SEPT3, for which a crystal structure employing a SEPT3 mutant (SEPT3_T282Y_) is available. However, the Cdc3-Cdc10 complex has both catalytically active and inactive subunits, different from SEPT7-SEPT3, where both subunits are catalytically active. Regarding solely this aspect, Cdc3-Cdc10 could instead be considered more similar to the human SEPT2-SEPT6 dimer which also forms a G-interface. Rosa et al., 2020, describe that the principal structural determinant of specificity at the SEPT2-SEPT6 interface involves switch I (also within the upper region) which makes a significant contribution to the total contact surface area between the two subunits. By contrast, the contact surface area involving switch I in the SEPT7-SEPT3_T282Y_ G-interface dimer is much smaller (Rosa et al., 2020). In the case of the Cdc3G-Cdc10G structure we observe a total contact area of 1,735 Å^2^, which is intermediate between those observed for SEPT2-SEPT6/8/11 (1,853 Å^2^) and SEPT7-SEPT3_T282Y_ (1,461 Å^2^). A significant contributor to the increase in the yeast complex when compared with its analog SEPT7-SEPT3_T282Y_ is indeed switch I, specifically that from Cdc10 which interacts with the trans-loop 1 of Cdc3 as observed in the structures of SEPT2-SEPT6/8/11. Ile66 in switch I of Cdc10, fits into a hydrophobic cavity formed by Tyr263, Leu264, Pro266 and Leu269 of Cdc3 in a manner similar to that observed for its homolog (Ala69) in SEPT2 in its complexes with SEPT6/8/11 (Fig. 5C).

On the other side of the interface lies the Switch I region of Cdc3, this is much larger than in other yeast septins and has a high density of negatively charged residues. It is disordered in the crystal structures and presumably projects into the solvent where its unusual size and charge appear to make it an ideal candidate for interacting with binding partners. An electrostatic surface representation of an AlphaFold2 predicted structure of the complete complex includes this region (Fig. S5) and highlights the negative charge density.

These observations indicate an unexpected similarity between the Cdc3-Cdc10 G-interface and that seen in the SEPT2-SEPT6/8/11 dimers rather than SEPT7-SEPT3_T282Y_, which is also supported by the values for the surface contact areas and types of nucleotides found buried in the interface. Thus, clustering Cdc3-Cdc10 together with human septins in an attempt to draw a coherent picture could be challenging and misleading since the yeast dimer G interface presents characteristics of both SEPT2-SEPT6 and SEPT7-SEPT3.

In the middle section of the interface two substantial changes are observed in Cdc3G-Cdc10G when compared to other known structures. First, the absence of the bifurcated salt bridge (Fig. 5D), and second, the substitution of an important and conserved His that interacts with the nucleotide across the interface (de Freitas Fernandes et al., 2022; Sirajuddin et al., 2009, 2007). The bifurcated salt bridge has been shown to be important in the stabilization of the G-interface in the human SEPT7 homodimer, since its absence in the *D. melanogaster* homologue (Pnut) is related to instability of the Pnut homodimer (de Freitas Fernandes et al., 2022). At the homodimeric G-interface in SEPT7 Glu58 and Lys173 from both subunits contribute to an approximately square planar arrangement of alternating charges in which each glutamic acid has two lysine neighbours and vice versa. This has been referred to as a bifurcated salt-bridge network. Although on one side of the Cdc3-Cdc10 interface Gln40 (Cdc10) and Lys265 (Cdc3) could potentially form a polar interaction, they are not observed to do so in the crystal structures and the lack of an acidic residue means that Lys265 is not compensated by a formal negative charge. On the other side of the interface, the equivalent residues are Ser158 and Pro127 preventing the formation of a network of interactions as seen in the SEPT7 homodimer (Fig. 5D) (de Freitas Fernandes et al., 2022; Rosa et al., 2020).

Normally, a histidine (His158 in SEPT2) of one subunit interacts directly with the α and/or β- phosphate of the nucleotide bound on the opposite side of the G-interface (Macedo et al., 2013; Rosa et al., 2020; Sirajuddin et al., 2009; Zent et al., 2011; Zeraik et al., 2014). In the yeast septins Cdc11, Cdc12 and Cdc3 the mutation of the homologous histidine by an alanine produces temperature sensitive cells and unobservable growth phenotypes. In addition, the His262Ala mutation in Cdc3 in the Cdc3-Cdc10 heterodimer, impairs the formation of stable heterodimers in solution (Sirajuddin et al., 2009). However, in SEPT2 the His158Ala mutation does not alter the catalytic activity of the protein, suggesting that His158 is more important in G-interface stabilization rather than nucleotide hydrolysis (Sirajuddin et al., 2009). Cdc10 shows a substitution of the conserved histidine for Lys155, a positive residue that directly interacts with the GTP γ-phosphate of Cdc3 (Fig. 5E). Although different in detail, this substitution maintains the interaction across the interface. This is presumably essential for the stability of the GTP bound to the complex and consequently the formation of an intact G-interface between Cdc3 and Cdc10.

Finally, in the lower section of the G-interface, there are novel stabilizing interactions in Cdc3- Cdc10 when compared to human SEPT7-SEPT3_T282Y_. The side chain of Arg254 found in the β- meander of Cdc10 forms a salt bridge with Asp293 at the beginning of helix α_4_ in Cdc3 (Fig. 5F). The latter interaction gives strength to the interface by the introduction of a charge-charge interaction and overall, the interface has a larger surface contact area, as quantified above. Importantly, the residues which participate in the additional salt bridge are absent from SEPT7 and SEPT3, where they are replaced by prolines in the homologous positions. In addition, we observed the conserved salt bridge between Arg251(Cdc10) and Glu295(Cdc3) related to the stabilization of the G-interface in septins (Fig. 5F) (de Freitas Fernandes et al., 2022). Molecular Dynamics simulations suggest that the lower region of the interface is where the strongest interactions reside as it is the last to break when opposite forces are applied to different monomers (de Freitas Fernandes et al., 2022; Grupp et al., 2023). The additional stability resulting from the salt bridge between Arg254(Cdc10) and Asp(293(Cdc3) may explain why yeast octamers are rarely seen to transform into hexamers due to the loss of the Cdc10 dimer whereas preparations of human octamers will frequently be contaminated by hexamers due to the loss of the SEPT3 subgroup member from the center of the particle (Sellin et al., 2011).

### Peculiarities of the Cdc3 structure: the GTP binding environment

Cdc3 is a catalytically inactive septin that has a lysine instead of a catalytic threonine (ThrSw1). Furthermore, it shows a substitution of a conserved serine/threonine in the *P-loop* for an aspartic acid at position 128. In SEPT2, a mutation in this conserved serine (Ser46, Fig. 6) by an aspartic acid reduces the catalytic activity of the protein and dramatically affects its affinity for the nucleotide (Sirajuddin et al., 2009). Several lines of evidence show that Cdc3 alone is unable to bind and hydrolyze nucleotides (both the full-length and the G-domain versions) (Baur et al., 2019). Nevertheless, when complexed to Cdc10, Cdc3 binds to GTP as was indirectly implied by the nucleotide release analysis and can be seen directly in the crystal structures. However, the structure of Cdc3 shows that it binds GTP in a slightly new arrangement to that seen previously (Fig. 6A).

**Figure 6.**
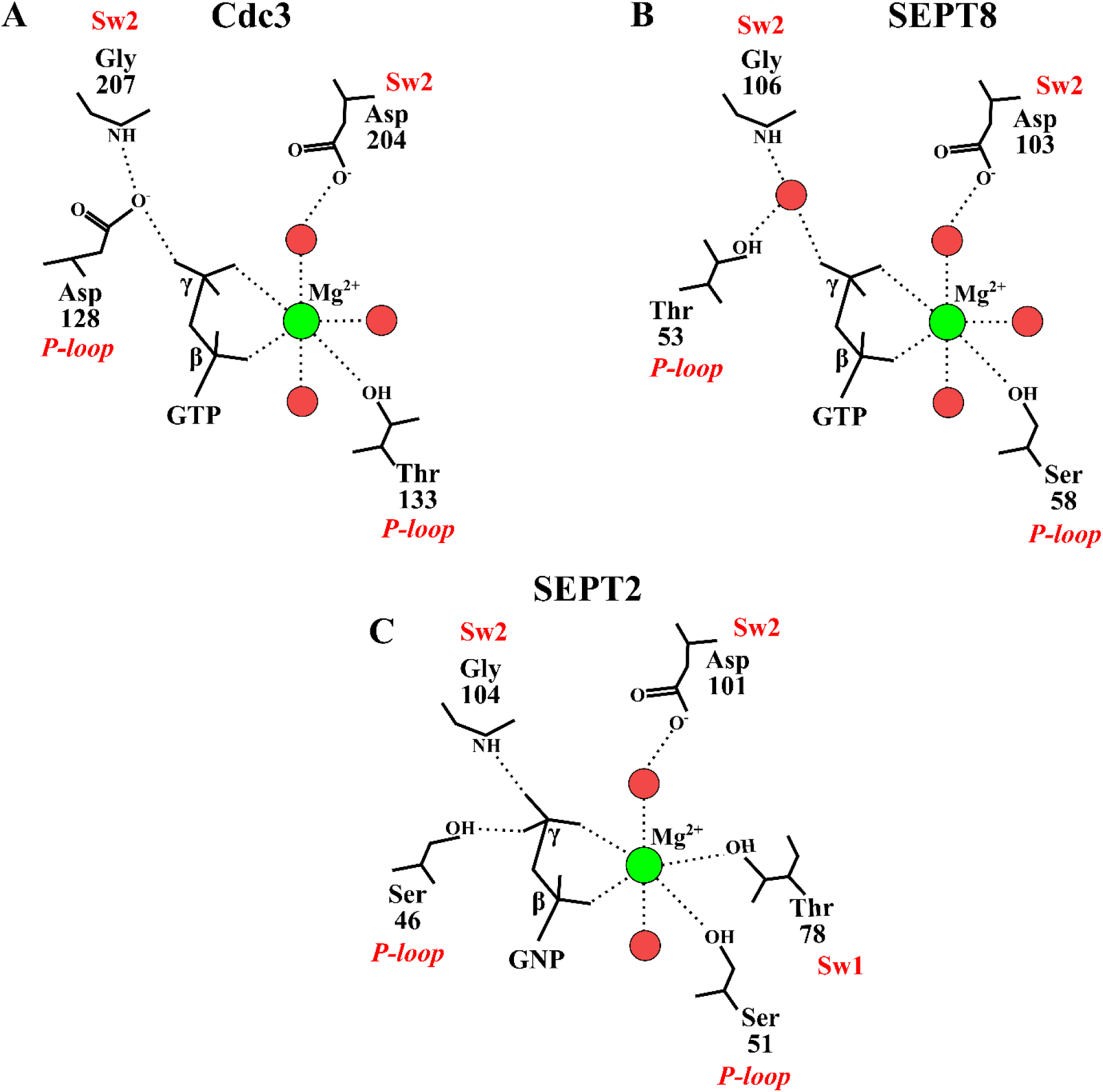
The nucleotide and Mg^2+^ coordination in Cdc3, SEPT8 (a catalytically inactive human septin), and SEPT2 bound to GNP (a non-hydrolyzed GTP homolog). A group of amino acids are conserved for the coordination of GTP and Mg^2+^, such as Thr/Ser in the P-loop that directly coordinates with magnesium, and Asp in switch II (Sw2). However, in Cdc3, Asp128 from the P-loop occupies the position of a conserved Thr/Ser that directly or indirectly binds to the γ-phosphate of GTP. The hexacoordination of magnesium is completed with a water molecule in Cdc3 and SEPT8 while in SEPT2 it is completed by the hydroxyl of the catalytic Thr78.

Comparing the GTP coordination in Cdc3 to that observed in SEPT2 bound to a non-hydrolyzed GTP analogue (GNP) and SEPT8 (a catalytically inactive human septin) subtle differences are observed. In SEPT2 the Mg^+2^ is coordinated by two water molecules, the β and γ phosphates from GTP and residues Ser51 (from the P-loop) and the catalytic Thr78 (from Switch I, ThrSwI) (Fig. 6C). One of the water molecules is stabilized by its interaction with Asp101 (from Switch 2), a conserved residue in septins. In SEPT8 and Cdc3, the Mg^2+^ lacks the direct coordination by the catalytic ThrSw1, which is impossible anyway as it is absent from both (Fig. 6A and 6B). Instead, it is replaced by a water molecule. The γ-phosphate in SEPT2 is coordinated directly by the P-loop Ser46 and the main chain of Gly104 from Switch 2 (Fig. 6C). However, with the substitution of the P-loop Ser/Thr for Asp128 in Cdc3, the coordination of Gly207 from Switch 2 with γ-phosphate is performed indirectly by the δ1 oxygen of Asp128. In SEPT8, this interaction is also indirect but via a water molecule (Fig, 6B). Therefore, in both inactive septins, besides the loss of the catalytic ThrSwI, there has been disruption to the constellation of interactions which comprise the classical “loaded spring” hydrolytic mechanism of small GTPases (Kozlova et al, 2022), albeit by different means.

### Comparison with the Cdc11 structure

Other than those reported here, the only other yeast septin structure which is currently available is that of Cdc11 (Brausemann et al., 2016). This is reported in its monomeric and apo form and presents some unusual features. These include the absolute values of the temperature factors (and their spatial distribution), identical values for R_work_ and R_free_ and significant structural differences within the Septin Unique Element (SUE) leading to a poor correlation between the structure and its electron density. A comparison between Cdc11 and the structures described here therefore seems warranted.

The structures of Cdc3 and Cdc10 present the classical septin fold including the SUE, a ∼50 residue segment at the C-terminus of the G-domain which forms the β-meander (β_a_, β_b_and β_c_, Cavini et al., 2021) and the helices α_5_ and α_6_. By contrast, in the structure of Cdc11 (Fig. 7) the β-meander is absent and a largely irregular structure occupies the region where one would expect to find α_5_. Furthermore, α_6_ is extended towards its N-terminus thereby eliminating the α_5_- α_6_ elbow. As described above, this is an important feature of the NC-interface and would be expected to be conserved in all septin structures, especially as it forms part of the SUE. It’s absence is surprising. On the other hand, it is only fair to mention that Cdc11 presents several novel features around the region of α_0_/PB1 indicating that the homodimeric NC interface which Cdc11 forms on octamer association into filaments, may not be totally conventional. Most noticeable, Cdc11 lacks the homologue of Gln21 which normally fits into the α_5_-α_6_ elbow (see above). This is replaced by a Leucine, which is unable to make equivalent hydrogen bonds. The Cdc11-Cdc11 NC-interface may yet reserve some surprises which have yet to be fully revealed and implies that a crystal structure where the appropriate NC contact is formed is rapidly becoming an imperative.

**Figure 7.**
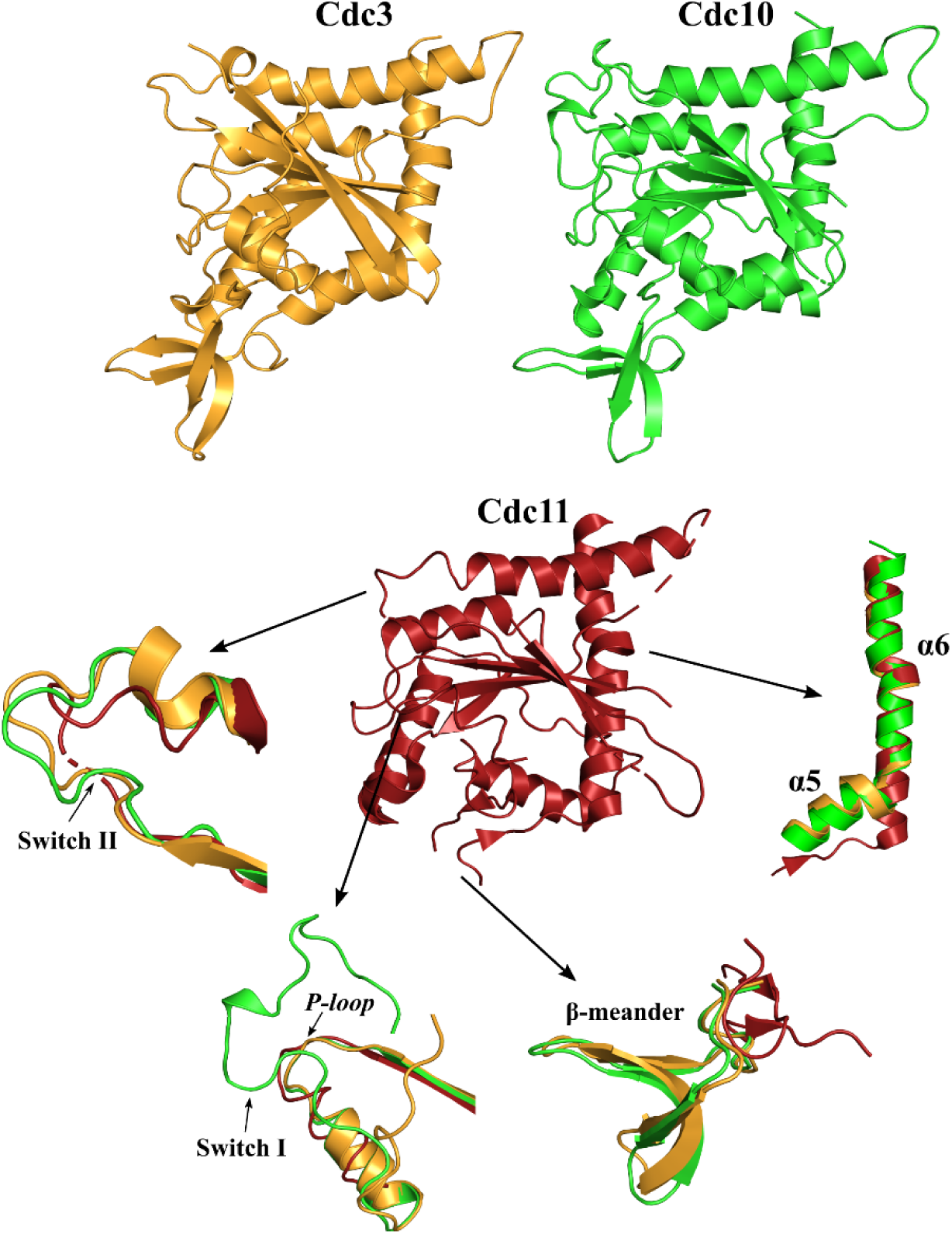
Comparison of the crystal structures of Cdc11, Cdc3, and Cdc10. The regions with the main differences between the structure of Cdc11 and Cdc3-Cdc10 are shown. Both switch I and switch II are more ordered in Cdc3-Cdc10. The β-meander, which was not modeled in Cdc11, has the canonical conformation in Cdc3-Cdc10. Finally, the absence of the α_5_-α_6_ elbow is notable in Cdc11.

Figure 7 compares some of the unusual features of Cdc11 with Cdc3 and Cdc10. These also include switches I and II which show greater levels of disorder than normally observed. This could potentially be explained by the absence of nucleotide and the lack of G-interface contacts within the crystal. Alternatively, it may be a further indication of anomalies in the Cdc11 structure itself. In summary, it would appear that the structure of Cdc11, including in its nucleotide-bound form, merits further investigation, especially as the AlphaFold2 prediction implies a structure with a conventional septin fold.

## Conclusions

Despite the considerable volume of molecular and cellular data which has accumulated on yeast septins over the last few decades, information on their structural biology has lagged sadly behind. This is particularly pertinent as it is not trivial to extrapolate structural conclusions from humans to yeast. For example, the catalytically active subunits are distributed differently in the two octamers and the individual septins have largely been classified into different groups (Shuman and Momany, 2022). The exception is Cdc10 and SEPT3. These share several common features and form NC-interface homodimers at the center of their respective protofilaments. Despite the differences, the structures provided here demonstrate many common features at both the G and NC interfaces. The Cdc3-Cdc10 pair occupies an analogous position to SEPT7-SEPT3 and retains some of its properties and yet has a G-interface which also shows some similarities with that observed for SEPT2-SEPT6 in the human particle. It has a contact surface area that is intermediate between those observed for SEPT7- SEPT3 and SEPT2-SEPT6 and involves the participation of switch I from Cdc10. The increased surface area compared with SEPT7-SEPT3, together with a contribution from Arg254 from the β-meander, probably imbues greater stability to the heterodimer, potentially explaining why yeast octamers appear to be more stable and homogenous when compared with those from humans which often dissociate into hexamers by losing the central SEPT3 (or SEPT9) dimer. For this to occur it is the smaller SEPT7-SEPT3 interface which must rupture.

The structure of the Cdc3G-Cdc10_Δ1-10_ provides a direct view of the central tetramer of the yeast protofilament. Much that we can now directly observe concerning the α_0_ helix of Cdc10 is consistent with mutation studies performed more than a decade ago and it’s special (possibly unique) properties concerning membrane interaction make it of particular interest (Bertin et al., 2010). As observed in the cryo-EM structure of the human SEPT2/6/7 complex (Mendonça et al., 2021), its basic residues are hidden within the interface where they largely participate in salt bridges. This seems to imply that a nucleotide-dependent squeezing mechanism exposing α_0_, as has been proposed for human septins, may also be applicable to yeast (Castro et al., 2020). Although there is no direct experimental evidence for this as yet, it appears to be consistent with the cross-talk between interfaces which has been described (Weems and McMurray, 2017).

No structure for the yeast octamer has yet been reported but it would be anticipated to show a considerable degree of flexibility, potentially greater than that seen for the human hexamer (Mendonça et al., 2021). Obtaining a high-resolution cryo-EM structure for it may prove challenging. The Cdc3G-Cdc10_Δ1-10_ structure provided here is an important piece of that puzzle and should aid towards achieving this goal as it represents half of the full particle and provides a resolution which should be appropriate for fitting into a cryo-EM map.

## Author contributions

RCG and APUA designed the research. RMS, GCRS and SAS carried out cell culture and sample preparation and conducted the biophysical characterization experiments by multi-angle light scattering. RMS carried out the nucleotide identification experiments. DAL and HMP carried out the crystallographic data collection and structure solution. RMS, DAL and RCG wrote the manuscript. RCG and APUA reviewed the manuscript.

## Formatting of funding sources

We gratefully acknowledge the support of the São Paulo Research Foundation (FAPESP) via grants 2014/15546-1 and 2020/02897-1 to R.C.G. and A.P.U.A., 2018/09687-2 to R.M.S., 2022/00125-7 to G.C.R.S and 2021/08158-9 to D.A.L.

## Acknowledgments

We are grateful to Derminda Isabel de Moraes and Andressa Alves Pinto for excellent technical support. This research used facilities at the Brazilian Synchrotron Light Laboratory (LNLS), part of the Brazilian Center for Research in Energy and Materials (CNPEM), a private non-profit organization under the supervision of the Brazilian Ministry for Science, Technology, and Innovation (MCTI). The Manacá beamline staff is acknowledged for assistance during the diffraction data collection experiments (project proposal 20231023).

## Data availability

The data that support the findings of this study are available in the Protein Data Bank: Cdc3G- Cdc10G with PDB code 8FWP and Cdc3G-Cdc10_Δ1-10_ with PDB code 8SGD.

## SUPPLEMENTAL INFORMATION

**Figure S1.**
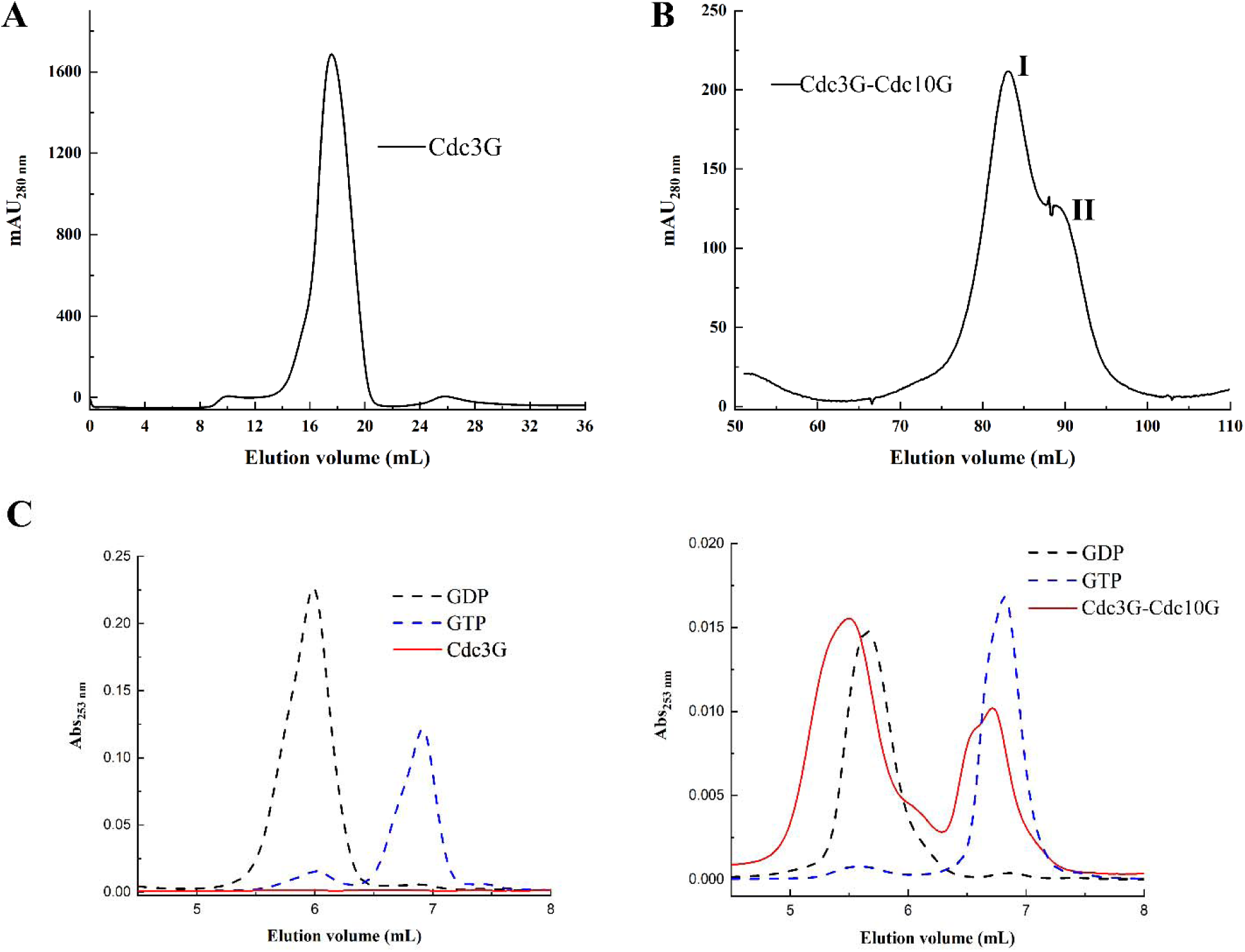
The size exclusion chromatography profile of A) Cdc3G and B) Cdc3G-Cdc10G shows a single oligomeric state for the former and two oligomeric states for the latter. SEC-MALS analysis reveals the molecular masses for each of the peaks (Fig. 1). C) Nucleotide identification from denatured protein shows apo Cdc3G in isolation but the presence of both GTP and GDP bound to Cdc3G-Cdc10G.

**Figure S2.**
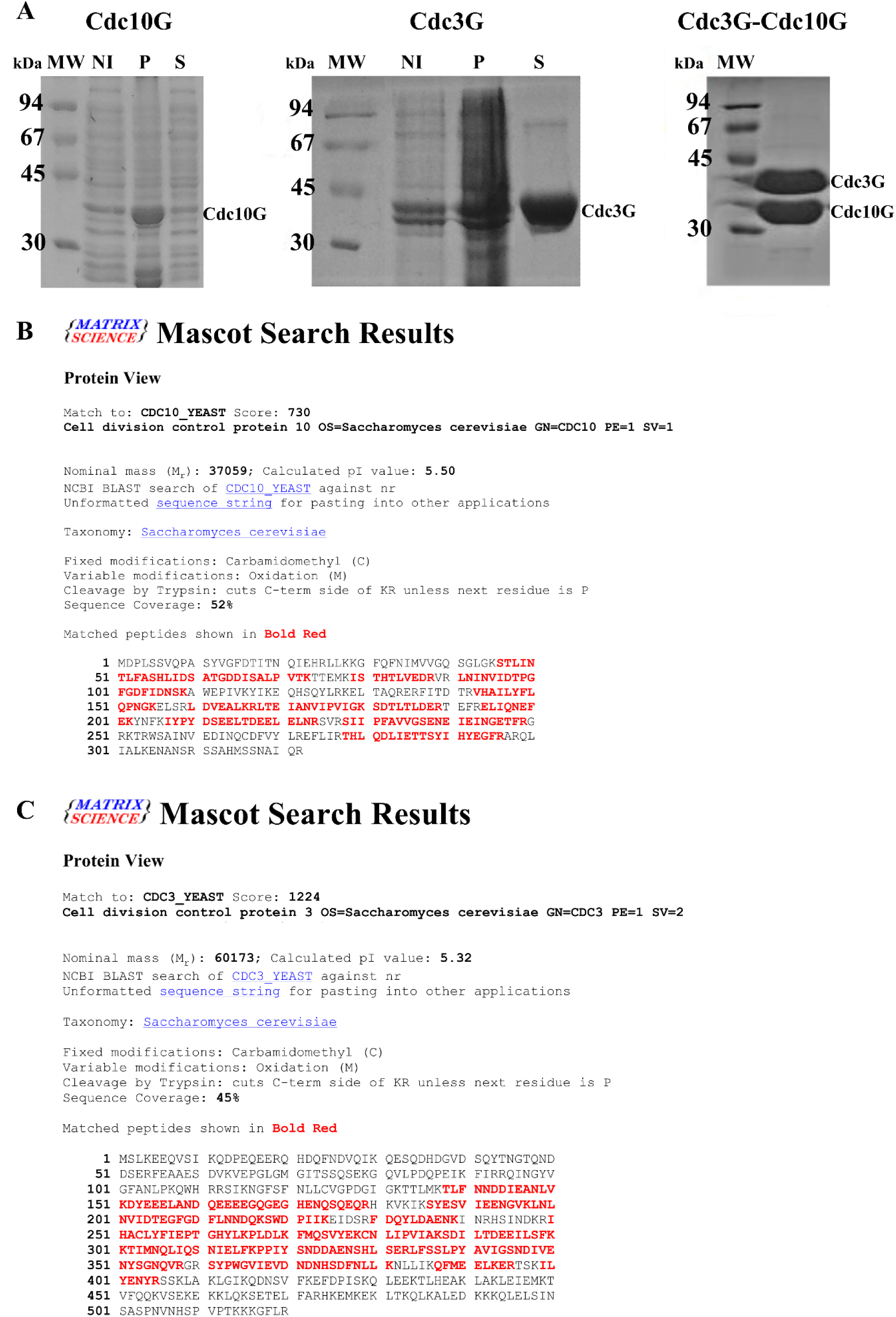
A) SDS-PAGE of recombinant Cdc10G, Cdc3G and the co-purification of Cdc3G- Cdc10G. When expressed separately, Cdc10G remains in the insoluble fraction (P) while Cdc3G remains largely in the soluble fraction (S). The bands are identified by their expected molecular masses to the right of the gel. NI=Not induced, P= Pellet and S=soluble. B) and C) show the peptides identified by mass spectrometry after trypsinization of the proteins extracted from the bands of the SDS-PAGE gels, corresponding to Cdc3G and Cdc10G.

**Figure S3.**
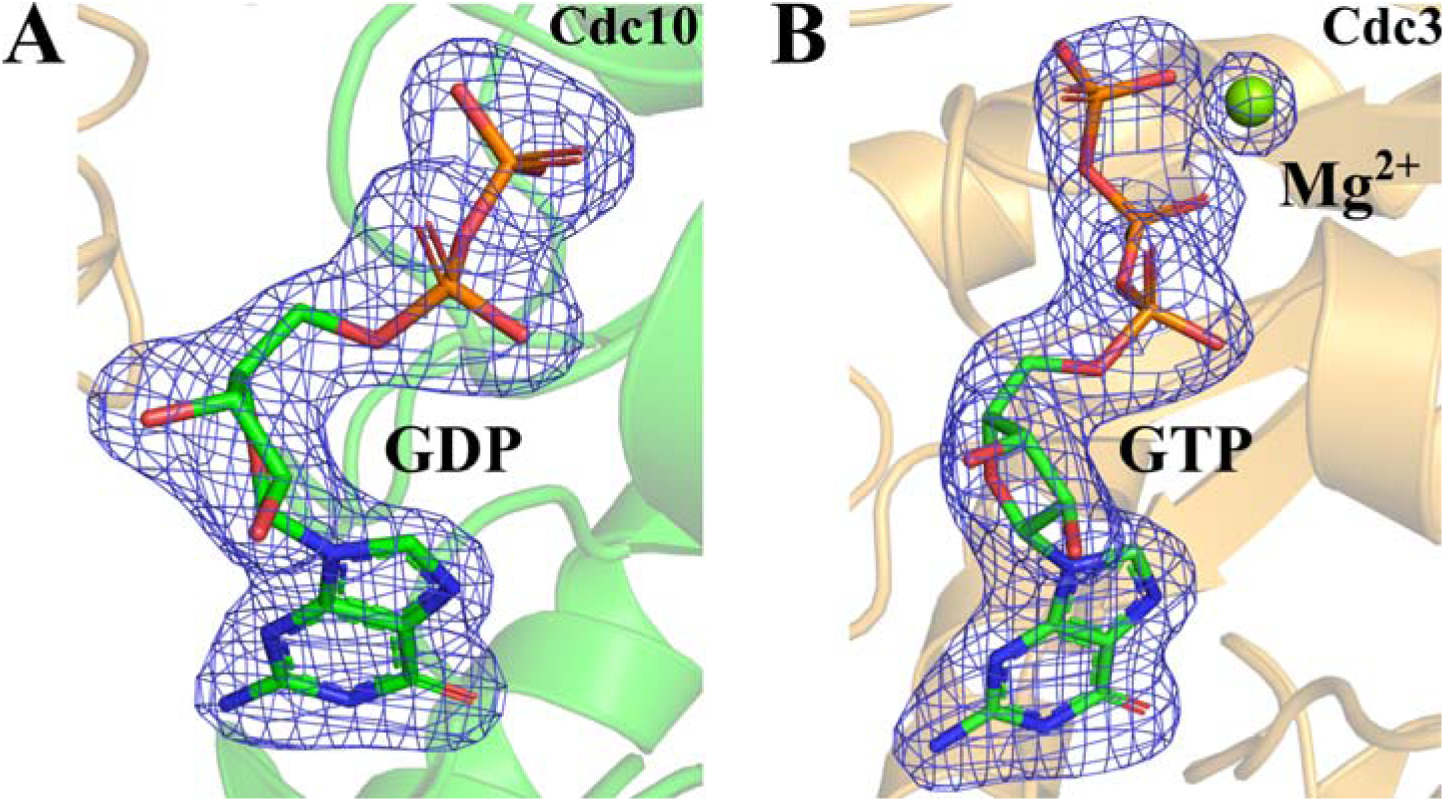
Polder map of GDP and GTP+Mg^2+^ bound to Cdc10 and Cdc3, respectively. Taken from the Cdc3G-Cdc10G complex.

**Figure S4.**
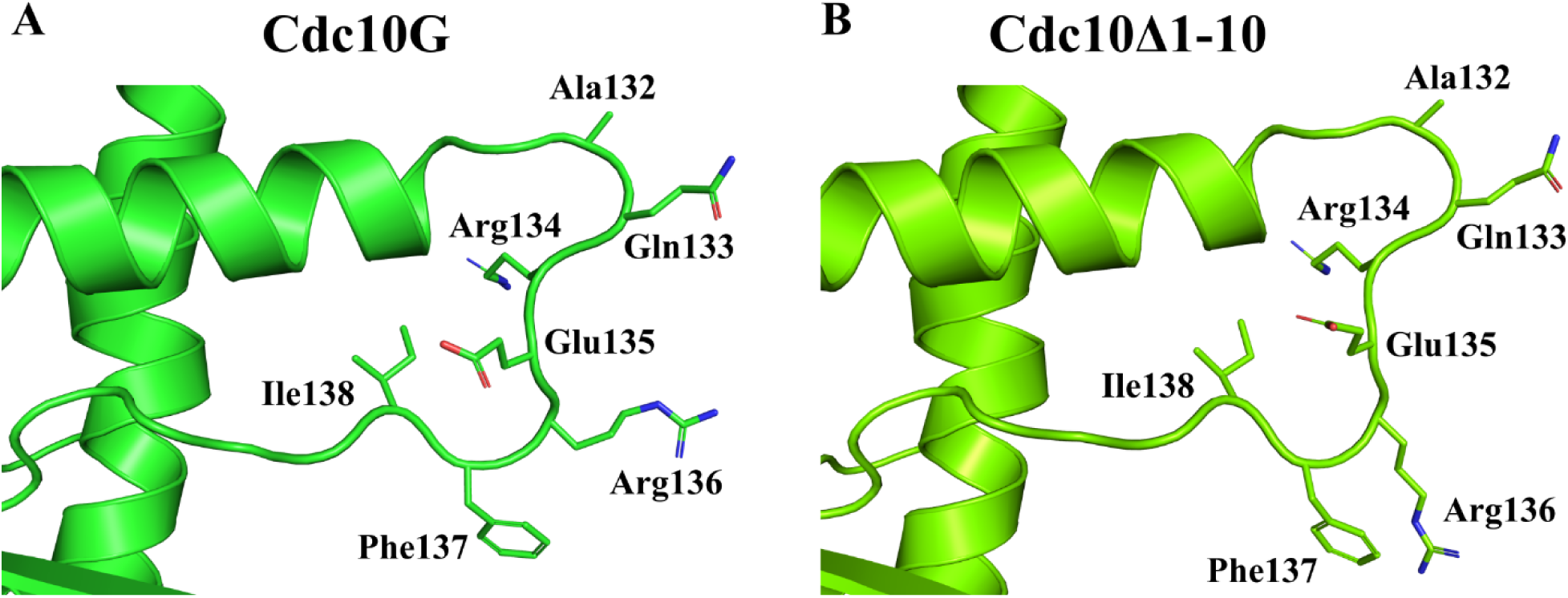
A) The α_2_-β_4_ loop (PB2) in Cdc10G and Cdc10_Δ1-10_. The orientation of the loop is conserved, however, Arg136 changes its rotamer due to the presence of the physiological NC interface in Cdc10_Δ1-10_, favoring the interaction with the PAR of the α_5’_-helix (not shown).

**Figure S5.**
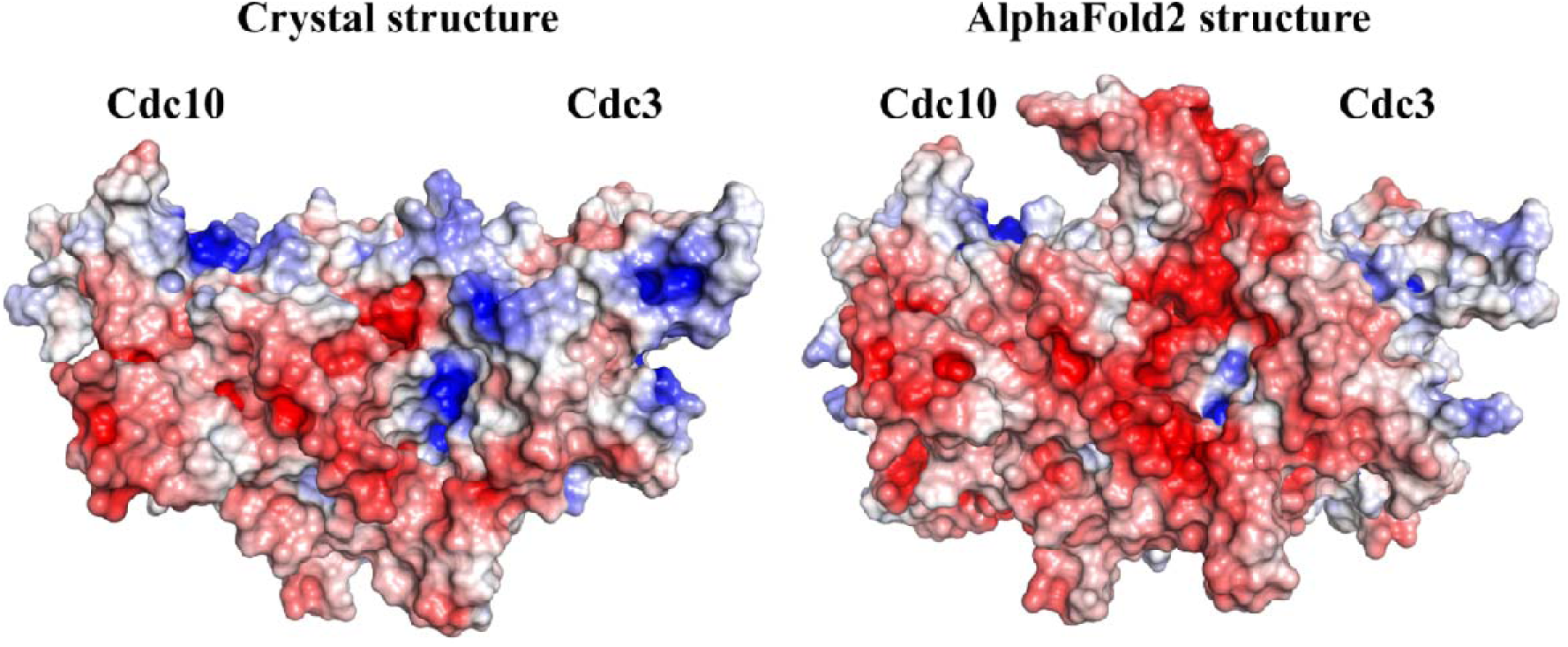
Surface representation colored according to electrostatic potential of the crystal structure of Cdc3G-Cdc10G and the AlphaFold2 model. The AlphaFold2 model shows a greater distribution of negative charges due to the presence of the complete switch I in Cdc3.

**Table S1.**
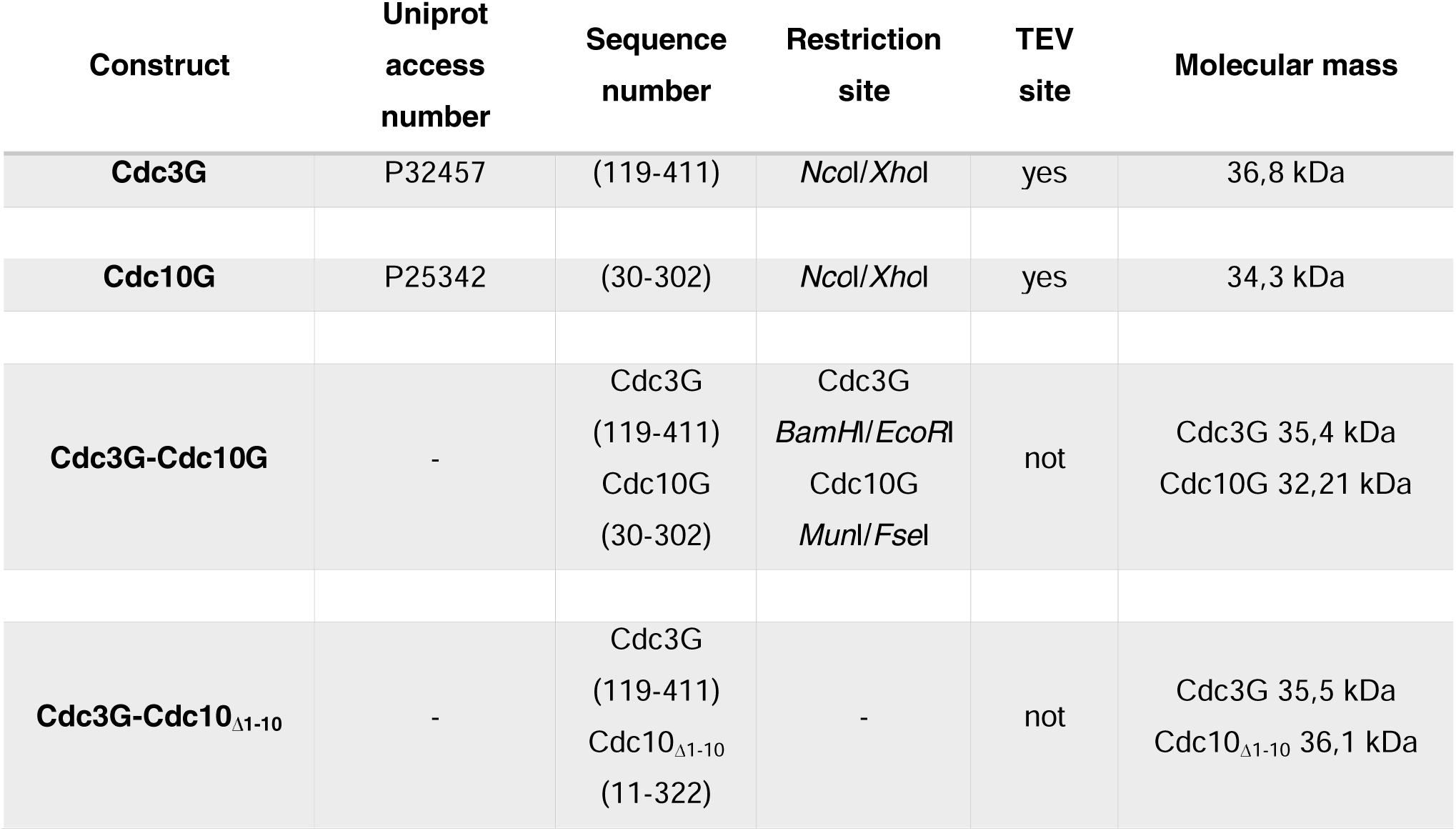
Details of the constructs used to produce the recombinant proteins.

| Construct | Uniprot<br>access<br>number | Sequence<br>number | Restriction<br>site | TEV<br>site | Molecular mass |
| --- | --- | --- | --- | --- | --- |
| **Cdc3G** | P32457 | (119-411) | *NcoI/XhoI* | yes | 36,8 kDa |
| **Cdc10G** | P25342 | (30-302) | *NcoI/XhoI* | yes | 34,3 kDa |
| **Cdc3G-Cdc10G** | - | Cdc3G<br>(119-411)<br>Cdc10G<br>(30-302) | Cdc3G<br>*BamHI/EcoRI*<br>Cdc10G<br>*MunI/FseI* | not | Cdc3G 35,4 kDa<br>Cdc10G 32,21 kDa |
| **Cdc3G-Cdc10<sub>Δ1-10</sub>** | - | Cdc3G<br>(119-411)<br>Cdc10<sub>Δ1-10</sub><br>(11-322) | - | not | Cdc3G 35,5 kDa<br>Cdc10<sub>Δ1-10</sub> 36,1 kDa |

**Table S2.**
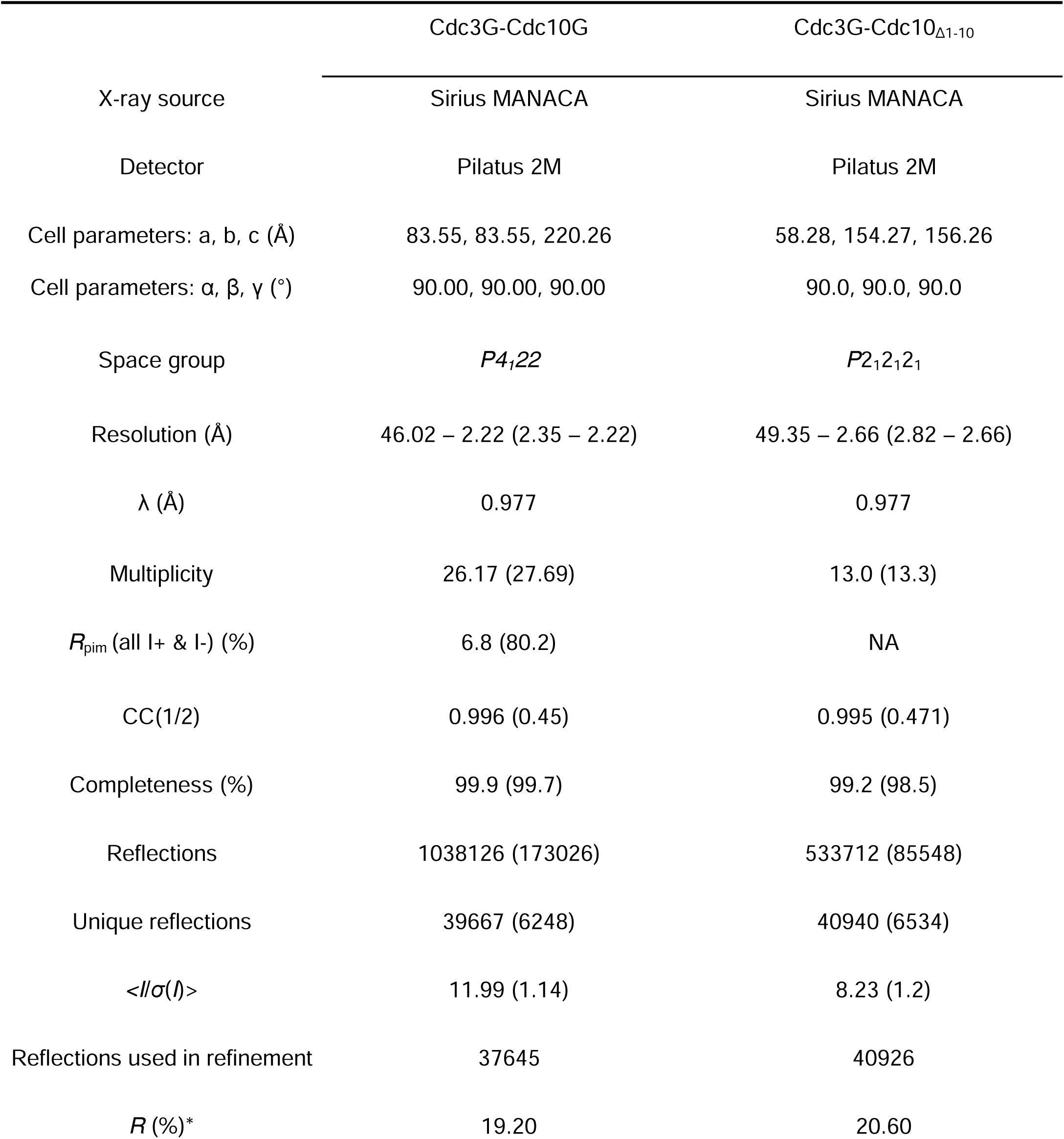
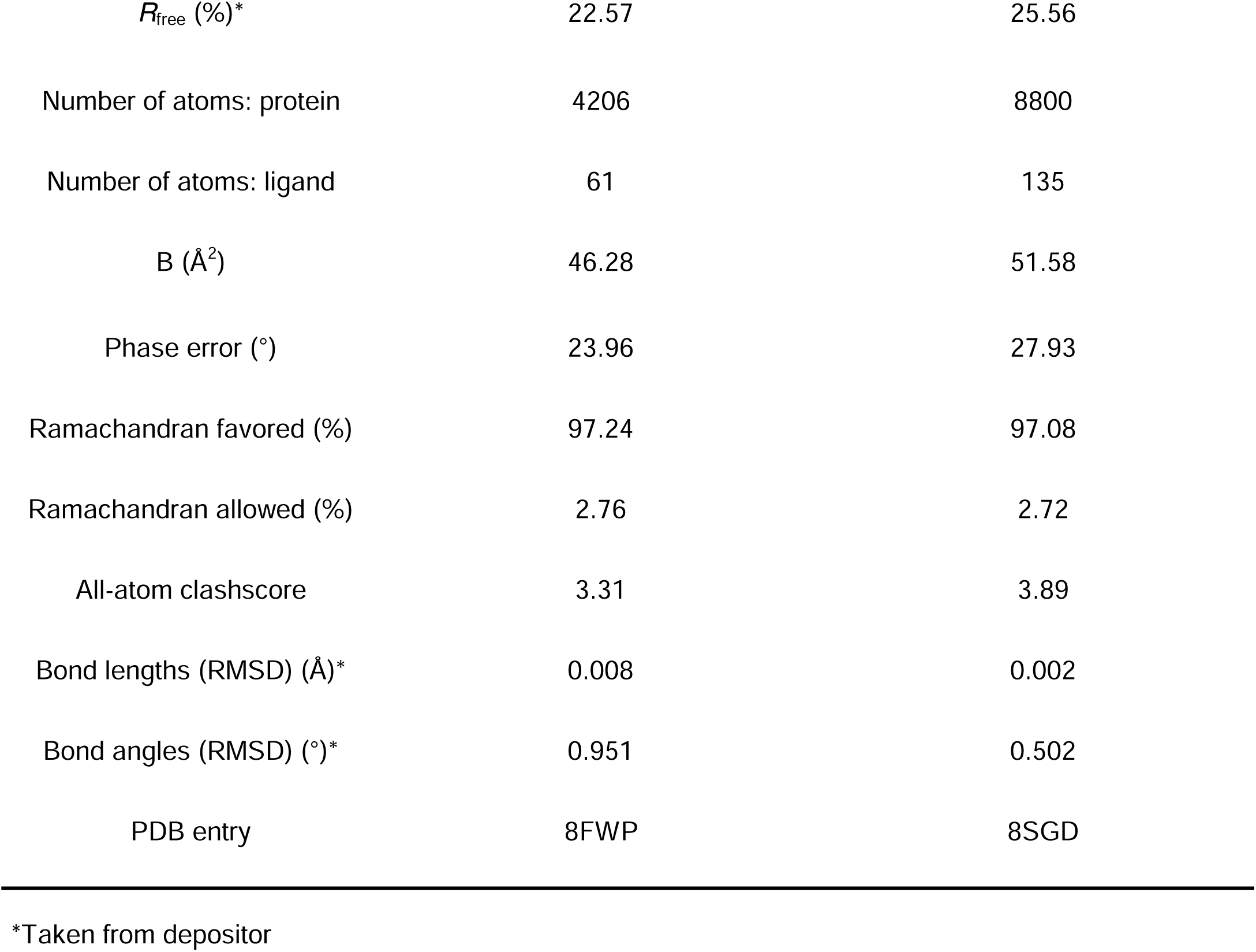
X-ray data collection and structure refinement statistics.

